# Time-dependent discrimination advantages for harmonic sounds suggest efficient coding for memory

**DOI:** 10.1101/2020.05.07.082511

**Authors:** Malinda J. McPherson, Josh H. McDermott

**Affiliations:** Department of Brain and Cognitive Sciences, MIT; Program in Speech and Hearing Biosciences and Technology, Harvard University; McGovern Institute for Brain Research, MIT; Center for Brains, Minds and Machines, MIT

## Abstract

Perceptual systems have finite memory resources and must store incoming signals in compressed formats. To explore whether representations of a sound’s pitch might derive from this need for compression, we compared discrimination of harmonic and inharmonic sounds across delays. In contrast to inharmonic spectra, harmonic spectra can be summarized, and thus compressed, using their fundamental frequency (f0). Participants heard two sounds and judged which was higher. Despite being comparable for sounds presented back-to-back, discrimination was better for harmonic than inharmonic stimuli when sounds were separated in time, implicating memory representations unique to harmonic sounds. Patterns of individual differences (correlations between thresholds in different conditions) indicated that listeners use different representations depending on the time delay between sounds, directly comparing the spectra of temporally adjacent sounds, but transitioning to comparing f0s across delays. The need to store sound in memory appears to determine reliance on f0-based pitch, and may explain its importance in music, in which listeners must extract relationships between notes separated in time.

## Introduction

Our sensory systems transduce information at high bandwidths, but have limited resources to hold this information in memory. In vision, short-term memory is believed to store schematic structure extracted from image intensities, e.g. object shape, or gist, that might be represented with fewer bits than the detailed patterns of intensity represented on the retina (1-4). For instance, at brief delays visual discrimination shows signs of being based on image intensities, believed to be represented in high-capacity but short-lasting sensory representations (5). By contrast, at longer delays more abstract (6, 7) or categorical (8) representations are implicated as the basis of short-term memory.

In other sensory modalities the situation is less clear. Audition, for instance, is argued to also make use of both a sensory trace and a short-term memory store (9, 10), but the representational characteristics of the memory store are not well characterized. There is evidence that memory for speech includes abstracted representations of phonetic features (11, 12) or categorical representations of phonemes themselves (13-15). Beyond speech, the differences between transient and persistent representations of sound remain unclear. This situation plausibly reflects a historical tendency within hearing research to favor simple stimuli, such as sinusoidal tones, for which there is not much to abstract or compress. Such stimuli have been used to characterize the decay characteristics of auditory memory (16-19), its vulnerability to interference (20, 21), and the possibility of distinct memory resources for different sound attributes (22-24), but otherwise place few constraints on the underlying representations.

Here we explore whether auditory perceptual representations could be explained in part by memory limitations. One widely proposed auditory representation is that of pitch (25, 26). Pitch is the perceptual property which enables sounds to be ordered from low to high (27), and is a salient characteristic of animal and human vocalizations, musical instrument notes, and some environmental sounds. Such sounds often contain harmonics, whose frequencies are integer multiples of a single fundamental frequency (f0; Figure 1a&b). Pitch is classically defined as the perceptual correlate of this f0, which is thought to be estimated from the harmonics in a sound even when the frequency component at the f0 is physically absent (Figure 1c). Despite the prevalence of this idea in textbook accounts of pitch, there is surprisingly little direct evidence that listeners utilize representations of a sound’s f0 when making pitch comparisons. For instance, discrimination of two harmonic sounds is normally envisioned to involve a comparison of estimates of the sounds’ f0s (25, 28). But if the frequencies of the sounds are altered to make them inharmonic (lacking a single f0; Figure 1d), discrimination remains accurate (28-31), even though there are no f0s to be compared. Such sounds do not have a pitch in the classical sense – one would not be able to consistently sing them back or otherwise match their pitch, for instance (32) – but listeners nonetheless hear a clear upward or downward change from one sound to the other, like that heard for harmonic sounds. This result is what would be expected if listeners were using the spectrum rather than the f0 (Figure 1e), e.g. by tracking frequency shifts between sounds (33). Although harmonic advantages are evident in some other tasks plausibly related to pitch perception (such as recognizing familiar melodies, or detecting out-of-key notes (31)), the cause of this task dependence remains unclear.

**Figure 1.**
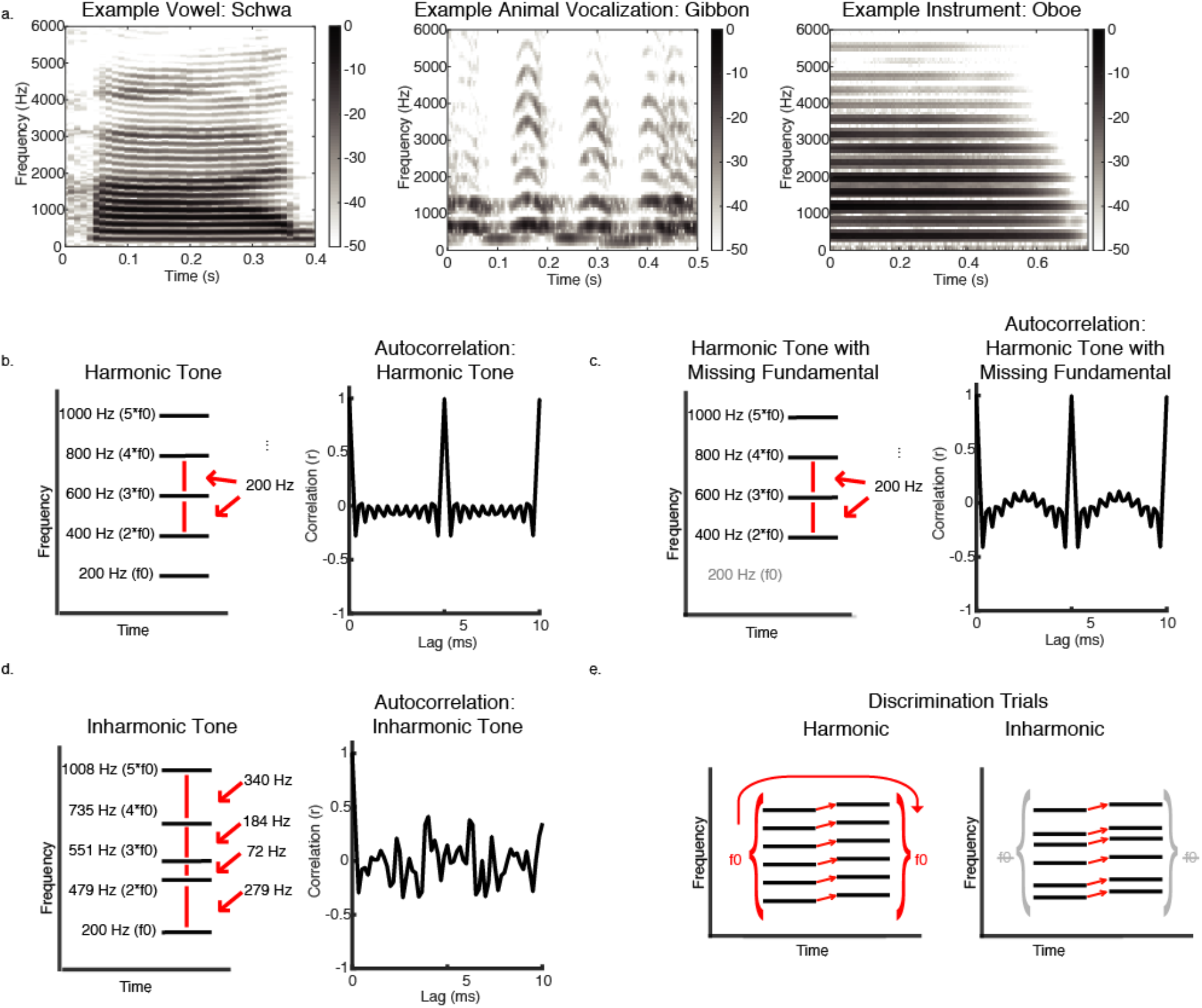
Example Harmonic and Inharmonic Sounds & Discrimination Trials. a. Example spectrograms for natural harmonic sounds, including a spoken vowel, the call of a gibbon monkey, and a note played on an oboe. The components of such sounds have frequencies that are multiples of an f0, and as a result are regularly spaced across the spectrum. b. Schematic spectrogram (left) of a harmonic tone with an f0 of 200 Hz along with its autocorrelation function (right). The autocorrelation has a value of 1 at a time lag corresponding to the period of the tone (1/f0 = 5 ms). c. Schematic spectrogram (left) of a harmonic tone (f0 of 200 Hz) missing its fundamental frequency, along with its autocorrelation function (right). The autocorrelation still has a value of 1 at a time lag of 5 ms, because the tone has the period of 200 Hz, even though this frequency is not present in its spectrum. d. Schematic spectrogram (left) of an inharmonic tone along with its autocorrelation function (right). The tone was generated by perturbing the frequencies of the harmonics of 200 Hz, such that the frequencies were not integer multiples of any single f0 in the range of audible pitch. Accordingly, the autocorrelation does not exhibit any strong peak. e. Schematic of trials in a discrimination task in which listeners must judge which of two tones is higher. For harmonic tones, listeners could compare f0 estimates for the two tones or follow the spectrum. The inharmonic tones cannot be summarized with f0s, but listeners could compare the spectra of the tones to determine which is higher.

In this paper we consider whether these characteristics of pitch perception could be explained by memory constraints. One reason to estimate a sound’s f0 might be that it provides an efficient summary of the spectrum: a harmonic sound contains many frequencies, but their values can all be predicted as integer multiples of the f0. This summary might not be needed if two sounds are presented back-to-back, as high-fidelity (but quickly-fading) sensory traces of the sounds could be compared. But it might become useful in situations where listeners are more dependent on a longer-lasting memory representation.

We explored this issue by measuring effects of time delay on discrimination. Our hypothesis was that time delays would cause discrimination to be based on short-term auditory memory representations (17, 34). We tested discrimination abilities with harmonic and inharmonic stimuli, varying the length of silent pauses between sounds being compared. We predicted that if listeners summarize harmonic sounds with a representation of their f0, then performance should be better for harmonic than for inharmonic stimuli. This prediction held across a variety of different types of sounds and task conditions, but only when sounds were separated in time. In addition, individual differences in performance across conditions indicate that listeners switch from representing the spectrum to representing the f0 depending on memory demands. Reliance on f0-based pitch thus appears to be driven in part by the need to store sound in memory.

## Results

### Experiment 1: Discriminating instrument notes with intervening silence

We began by measuring discrimination with and without an intervening delay between notes (Figure 2a), using recordings of real instruments that were resynthesized to be either harmonic or inharmonic (Figure 2b). We used real instrument sounds to maximize ecological relevance. Here and in subsequent experiments, sounds were made inharmonic by adding a random frequency ‘jitter’ to each successive harmonic; each harmonic could be jittered in frequency by up to 50% of the original f0 of the tone (jitter values were selected from a uniform distribution subject to constraints on the minimum spacing between adjacent frequencies). Making sounds inharmonic renders them inconsistent with any f0, such that the frequencies cannot be summarized by a single f0. Whereas the autocorrelation function of a harmonic tone shows a strong peak at the period of the f0 (Figure 1b-c), that for an inharmonic tone does not (Figure 1d). The same pattern of random ‘jitter’ was added to each of the two notes in a trial. There was thus a direct correspondence between the frequencies of the first and second notes even though the inharmonic stimuli lacked an f0 (Figure 1e).

**Figure 2.**
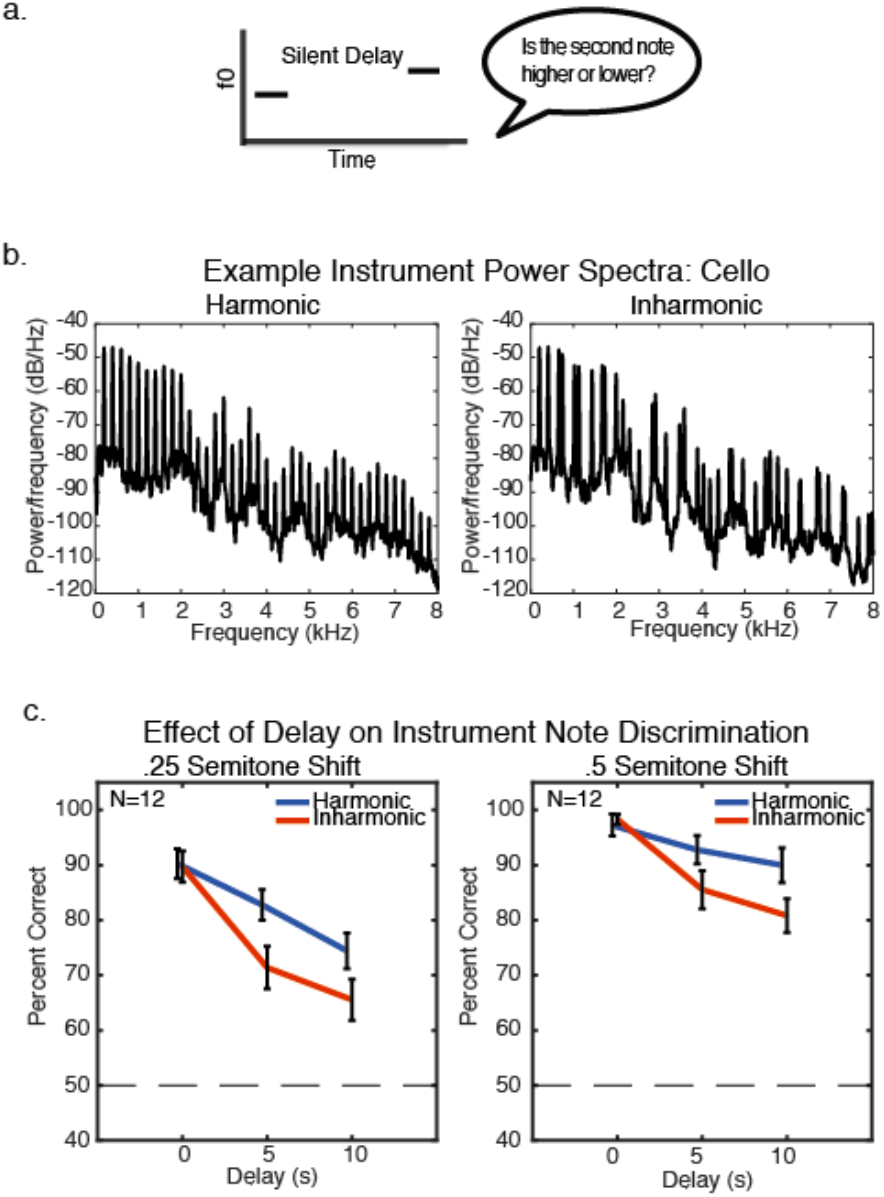
Experiment 1 – Harmonic advantage when discriminating instrument notes across a delay. a. Schematic of trial structure for Experiment 1. During each trial, participants heard two notes played by the same instrument and judged whether the second note was higher or lower than the first note. Notes were separated by a delay of 0, 5, or 10 seconds. b. Power spectra of example harmonic and inharmonic (with frequencies jittered) notes from a cello (the fundamental frequency of the harmonic note is 200 Hz in this example). c. Results of Experiment 1 plotted separately for the two difficulty levels that were used. Error bars show standard error of the mean.

Participants heard two notes played by the same instrument (randomly selected on each trial from the set of cello, baritone saxophone, ukulele, pipe organ, and oboe, with the instruments each appearing an equal number of times within a given condition). The two notes were separated by 0, 5, or 10 seconds of silence. Notes always differed by either a quarter of a semitone (approximately 1.5% difference between the note f0s) or a half semitone (approximately 3% difference between the note f0s). Participants judged whether the second note was higher or lower than the first.

We found modest decreases in performance for harmonic stimuli as the delay increased (significant main effect of delay for both .25 semitone (F(2,22)=25.19, p<.001, η_p_^2^=.70, Figure 2c), and .5 semitone conditions (F(2,22)=5.31, p=.01, η_p_^2^=.33)). This decrease in performance is consistent with previous studies examining memory for complex tones (21, 35). We observed a more pronounced decrease in performance for inharmonic stimuli, with worse performance than for harmonic stimuli at both delays (5 seconds: t(11)=4.48, p<.001 for .25 semitone trials, t(11)=2.12, p=.057 for .5 semitone trials; 10 seconds: t(11)=3.64, p=.004 for .25 semitone trials, t(11)=4.07, p=.002 for .5 semitone trials) despite indistinguishable performance for harmonic and inharmonic sounds without a delay (t(11)=0.43, p=.67 for .25 semitone trials, t(11)=-0.77, p=.46 for .5 semitone trials). These differences produced a significant interaction between the effect of delay and that of harmonicity (F(2, 22)=7.66, p=.003, η_p_^2^=.41 for .25 semitone trials, F(2, 22)=3.77, p=.04, η_p_^2^=.26 for .5 semitone trials). This effect was similar for musicians and nonmusicians (Supplementary Figure 1). Averaging across the two difficulty conditions we found no main effect of musicianship (F(1,10)=3.73, p=.08, η_p_^2^=.27), no interaction between musicianship, harmonicity and delay length (F(2,20)=0.58, p=.57, η_p_^2^=.06), and the interaction between delay and harmonicity was significant in non-musicians alone (F(2,10)=6.48, p=.02, η_p_^2^=.56). This result suggests that the harmonic advantage is not dependent on extensive musical training. Overall, the results of Experiment 1 are consistent with the idea that a sound’s spectrum can mediate discrimination over short time intervals (potentially via a sensory trace of the spectrum), but that memory over longer periods relies on a representation of f0.

### Experiment 2: Discriminating synthetic tones with intervening silence

We replicated and extended the results of Experiment 1 using synthetic tones, the acoustic features of which can be more precisely manipulated. We generated complex tones that were either harmonic or inharmonic, applying fixed bandpass filters to all tones in order to minimize changes in the center of mass of the tones that could otherwise be used to perform the task (Figure 3a). The first audible harmonic of these tones was generally the 4^th^ (though it could be the 3^rd^ or 5^th^, depending on the f0 or jitter pattern). To gauge the robustness of the effect across difficulty levels, we again used two f0 differences (.25 and .5 semitones).

**Figure 3.**
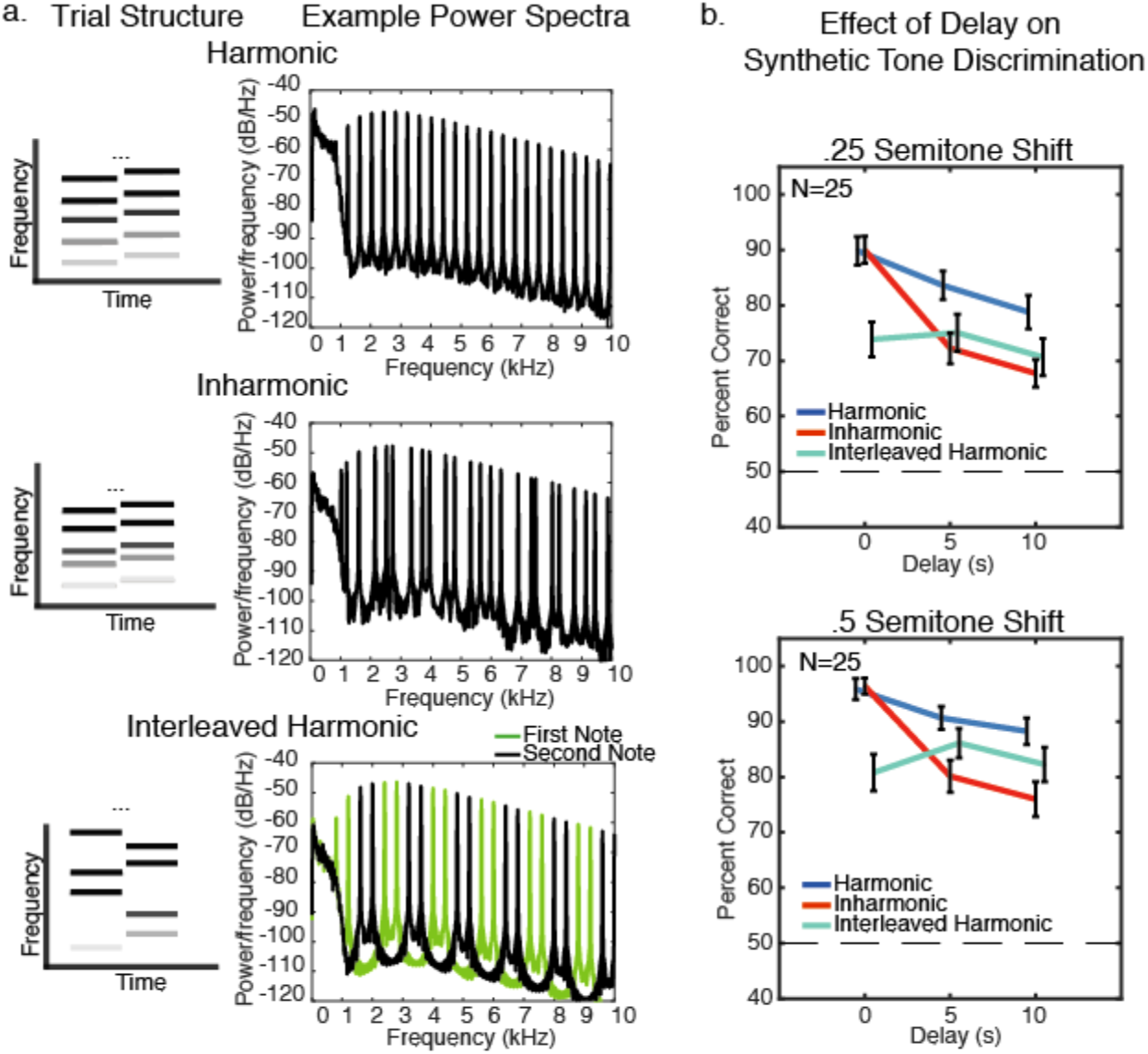
Experiment 2 – Harmonic advantage when discriminating synthetic tones across a delay. a. Schematic of stimuli and trial structure in Experiment 2. Left: During each trial, participants heard two tones and judged whether the second was higher or lower than the first. Tones were separated by a delay of 0, 5, or 10 seconds. Right: Power spectra of 400 Hz tones for Harmonic, Inharmonic, and Interleaved Harmonic conditions. b. Results of Experiment 2, plotted separately for the two difficulty levels. Error bars show standard error of the mean.

To further probe the effect of delay on representations of f0, we included a third condition (‘Interleaved Harmonic’) where each of the two tones on a trial contained half of the harmonic series (Figure 3a). One tone always contained harmonics [1, 4, 5, 8, 9, etc.], and the other always contained harmonics [2, 3, 6, 7, 10, etc.] (28). The order of the two sets of harmonics was randomized across trials. This selection of harmonics eliminates common harmonics between tones. While in the Harmonic and Inharmonic conditions listeners could use the correspondence of the individual harmonics to compare the tones, in the Interleaved Harmonic condition there is no direct correspondence between harmonics present in the first and second notes, such that the task can only be performed by estimating and comparing the f0s of the tones. By applying the same bandpass filter to each Interleaved Harmonic tone we sought to minimize timbral differences that are known to impair f0 discrimination (36), though the timbre nonetheless changed somewhat from note-to-note, which one might expect would impair performance to some extent. Masking noise was included in all conditions to prevent distortion products from being audible, which might otherwise be used to perform the task. The combination of the bandpass filter and the masking noise was also sufficient to prevent the frequency component at the f0 from being used to perform the task.

As in Experiment 1, harmonic and inharmonic tones were similarly discriminable with no delay (Figure 3c; Z=.55 p=.58 for .25 semitone trials, Z=0.66, p=.51 for .5 semitone trials, Wilcoxon signed-rank test). However, performance with inharmonic tones was again significantly worse than performance with harmonic tones with a delay (5-second delay: Z=3.49, p<.001 for .25 semitones, Z=3.71, p<.001 for .5 semitones; 10-second delay: Z=3.46 p<.001 for .25 semitones, Z=4.01, p<.001 for .5 semitones), yielding interactions between the effect of delay and harmonicity (F(2,48)=10.71, p<.001, η_p_^2^=.31 for .25 semitones, F(2,48)=26.17, p<.001, η_p_^2^=.52 for .5 semitones, p values calculated via bootstrap because data were non-normal).

Performance without a delay was worse for interleaved-harmonic tones than for the regular harmonic and inharmonic tones, but unlike in those other two conditions, Interleaved Harmonic performance did not deteriorate significantly over time (Figure 3b). There was no main effect of delay for either .25 semitones (F(2,48)=1.23, p=.30, η_p_^2^=.05) or .5 semitones (F(2,48)=2.56, p=.09, η_p_^2^=.10), in contrast to the significant main effects of delay for Harmonic and Inharmonic conditions at both difficulty levels (p<.001 in all cases). This pattern of results is consistent with the idea that there are two representations that listeners could use for discrimination: a sensory trace of the spectrum which decays quickly over time, and a representation of the f0 which is better retained over a delay. The spectrum can be used in the Harmonic and Inharmonic conditions, but not in the Interleaved Harmonic condition.

As in Experiment 1, the effects were qualitatively similar for musicians and non-musicians (Supplementary Figure 2). Although there was a significant main effect of musicianship (F(1,23)=10.28, p<.001, η_p_^2^=.99), the interaction between the effects of delay and harmonicity was significant in both musicians (F(2,28)=20.44, p<.001, η_p_^2^=.59) and non-musicians (F(2,18)=11.99, p<.001, η_p_^2^=.57), and there was no interaction between musicianship, stimulus type (Harmonic, Inharmonic, Interleaved Harmonic), and delay length (F(4,92)=0.19, p=.98, η_p_^2^=.01).

### Experiment 3: Discriminating synthetic tones with a consistent inharmonic spectrum

The purpose of Experiment 3 was to examine the effects of the specific inharmonicity manipulation used in Experiments 1 and 2. In Experiments 1 and 2, we used a different inharmonic jitter pattern for each trial. In principle, listeners might be able to learn a spectral pattern if it repeats across trials (37, 38), such that the harmonic advantage found in Experiments 1 and 2 might not have been due to harmonicity per se, but rather to the fact that the harmonic pattern occurred repeatedly whereas the inharmonic pattern did not. In Experiment 3 we compared performance in a condition where the jitter pattern was held constant across trials (‘Inharmonic-Fixed’) vs. when it was altered every trial (though the same for the two notes of a trial), as in the previous experiments (‘Inharmonic’).

Unlike the first two experiments, we used adaptive procedures to measure discrimination thresholds. Adaptive threshold measurements avoided difficulties associated with choosing stimulus differences appropriate for the anticipated range of performance across the conditions. Participants completed 3-down-1-up two-alternative-forced-choice (‘is the second tone higher or lower than the first’) adaptive ‘runs’, each of which produced a single threshold measurement. Tones were either separated by no delay, a 1-second delay, or a 3-second delay (Figure 4a). There were three stimulus conditions, separated into three blocks: Harmonic (as in Experiment 2), Inharmonic, where the jitter changed for every trial (as in Experiment 2), and Inharmonic-Fixed, where a single random jitter pattern (chosen independently for each participant) was used across the entire block of adaptive runs. Participants completed four adaptive runs for each delay and stimulus pair. Delay conditions were intermixed within the Harmonic, Inharmonic and Inharmonic-Fixed stimulus condition blocks, resulting in 12 adaptive runs per block (Figure 4b), with approximately 60 trials per run on average. The order of these three blocks was randomized across participants. This design does not preclude the possibility that participants might learn a repeating inharmonic pattern given even more exposure to it, but it puts the inharmonic and harmonic tones on equal footing, testing whether the harmonic advantage might be due to relatively short-term learning of the consistent spectral pattern provided by harmonic tones.

**Figure 4.**
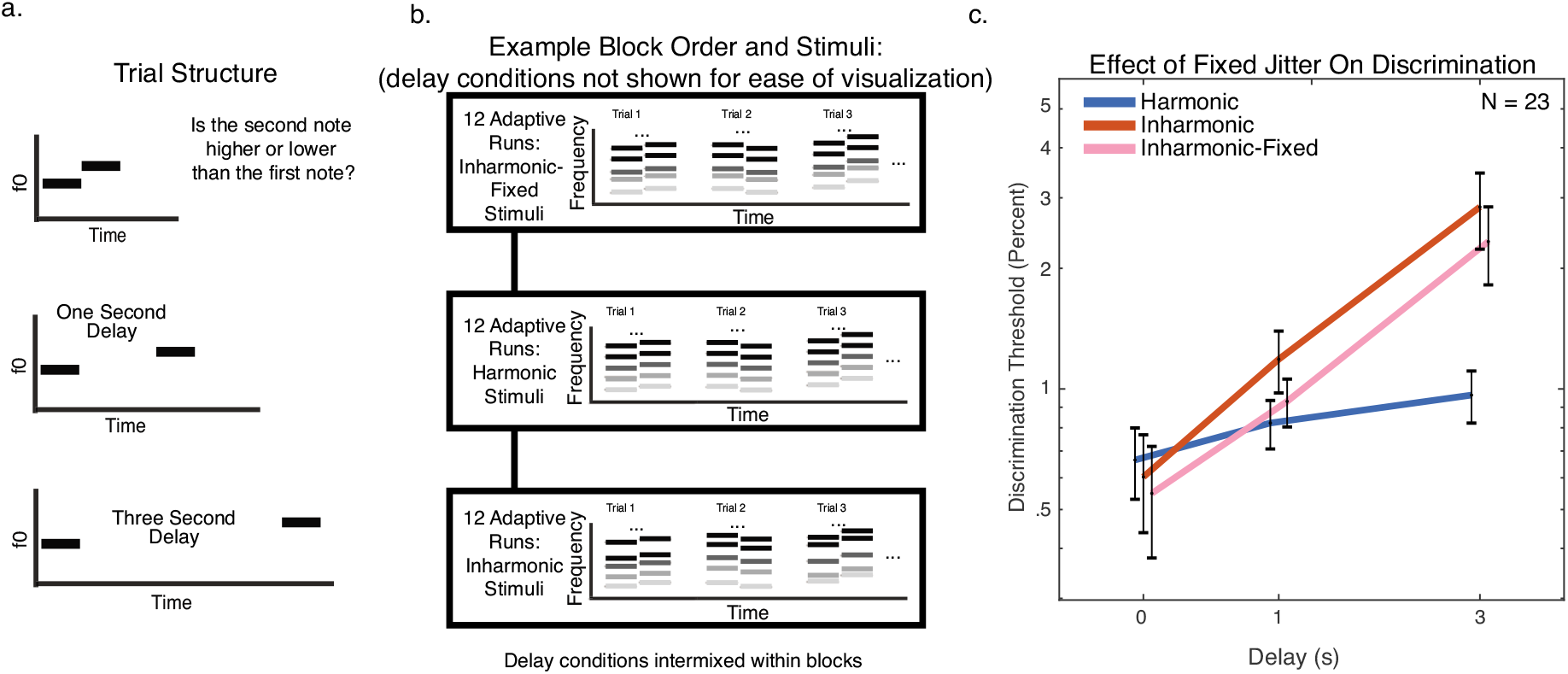
Experiment 3 – Harmonic advantage persists for consistent inharmonic jitter pattern. a. Schematic of trial structure for Experiment 3. Task was identical to that of Experiments 1 and 2, but with delay durations of 0, 1 and 3 seconds, and adaptive threshold measurements rather than method of constant stimuli. b. Example block order for Experiment 3, in which the 12 adaptive runs for each condition were presented within a contiguous block. The beginning of an example run is shown schematically for each type of condition. Stimulus conditions (Harmonic, Inharmonic, Inharmonic-Fixed) were blocked. Note the difference between the Inharmonic-Fixed condition, in which the same jitter pattern was used across all trials within a block, and the Inharmonic condition, in which the jitter pattern was different on every trial. Delay conditions were intermixed within each block. c. Results of Experiment 3. Error bars show within-subject standard error of the mean.

As shown in Figure 4c, thresholds were similar for all conditions with no delay (no significant difference between any of the conditions (Z<0.94, p>.34 for all pairwise comparisons, Wilcoxon signed-rank test, used because the distribution of thresholds was non-normal). These discrimination thresholds were comparable to previous measurements for stimuli with resolved harmonics (39, 40). However, thresholds were slightly elevated (worse) for both the Inharmonic and Inharmonic-Fixed conditions with a 1 second delay (Inharmonic: Z=2.40, p=.016; Inharmonic-Fixed: Z=1.92, p=.055), and were much worse in both conditions with a 3-second delay (Inharmonic: Z=3.41, p<.001; Inharmonic-Fixed: Z=2.71, p=.007). There was no significant effect of the type of inharmonicity (F(1,22)=0.94, p=.34, η_p_^2^=.04, comparing Inharmonic vs. Inharmonic-Fixed conditions), providing no evidence that participants learn to use specific jitter patterns over the course of a typical experiment duration. We again observed significant interactions between the effects of delay and stimulus type (Harmonic, Inharmonic, and Inharmonic-Fixed conditions) in both musicians (F(4,44)=3.85, p=.009, η_p_^2^=.26) and non-musicians (F(4,40)=3.04, p=.028, η_p_^2^=.23), and no interaction between musicianship, stimulus type, and delay length (F(4,84)=1.05, p=.07, η_p_^2^=.05, see Supplementary Figure 3).

These results indicate that the harmonic advantage cannot be explained by the consistency of the harmonic spectral pattern across trials, as making the inharmonic spectral pattern consistent did not reduce the effect. Given this result, in subsequent experiments we opted to use Inharmonic rather than Inharmonic-Fixed stimuli, to avoid the possibility that the results might otherwise be biased by the choice of a particular jitter pattern.

### Experiment 4: Discriminating synthetic tones with a longer inter-trial interval

To assess whether the inharmonic deficit could somehow reflect interference from successive trials (41) rather than the decay of memory during the inter-stimulus delay, we replicated a subset of the conditions from Experiment 3 using a longer inter-trial interval. We included only the Harmonic and Inharmonic conditions, with and without a 3 second delay between notes. For each condition participants completed four adaptive threshold measurements without any enforced delay between trials, and four adaptive thresholds where we imposed a 4 second inter-trial interval (such that the delay between trials would always be at least 1 second longer than the delay between notes of the same trial). This experiment was run online because the lab was temporarily closed due to the COVID-19 virus.

The interaction between within-trial delay (0 vs. 3 seconds) and stimulus type (Harmonic vs. Inharmonic) was present both with and without the longer inter-trial interval (with: F(1,37)=4.92, p=.03, η_p_^2^=.12; without: F(1,37)=12.34, p=.001, η_p_^2^=.25). In addition, we found no significant interaction between inter-trial interval, within-trial delay, and harmonic vs. inharmonic stimuli (F(1,37)=1.78, p=.19, η_p_^2^=.05, Supplementary Figure 4). This result suggests that the inharmonic deficit is due to difficulties retaining a representation of the tones during the delay period, rather than some sort of interference from preceding stimuli.

### Experiments 5: One-shot discrimination with a longer intervening delay

Experiments 1-4 leave open the possibility that listeners might use active rehearsal of the stimuli (singing to themselves, for instance) to perform the task over delays. Although prior results suggest that active rehearsal does not obviously aid discrimination of tones over delays (17, 42), it could in principle explain the harmonic advantage on the assumption that it is more difficult to rehearse an inharmonic stimulus, and so it seemed important to address. To assess whether the harmonic advantage reflects rehearsal, we ran a ‘one-shot’ online experiment with much longer delay times, during which participants filled out a demographic survey. We assumed this unrelated task would prevent them from actively rehearsing the heard tone. This experiment was run online to recruit the large number of participants needed to obtain sufficient power. We have previously found that online participants can perform about as well as in-lab participants (38, 43) provided basic steps are taken both to maximize the chances of reasonable sound presentation by testing for earphone/headphone use (44), and to ensure compliance with instructions, either by providing training or by removing poorly performing participants using hypothesis-neutral screening procedures.

Each participant completed only two trials in the main experiment. One trial had no delay between notes, as in the 0 s delay conditions of the previous experiments. During the other trial participants heard one tone, then were redirected to a short demographic survey, and then heard the second tone (Figure 5a). The order of the two trials was randomized across participants. For each participant, both trials contained the same type of tone, randomly assigned. The tones were either harmonic, inharmonic, or interleaved-harmonic (each identical to the tones used in Experiment 2). The two test tones always differed in f0 by a semitone. The discrimination task was described to participants at the start of the experiment, such that participants knew they should try to remember the first tone before the survey and that they would be asked to compare it to a second tone heard after the survey. To ensure task comprehension, participants completed 10 practice trials with feedback (without a delay, with an f0 difference of a semitone). These practice trials were always with same types of tones the participant would hear during the main experiment (for instance, if participants heard inharmonic stimuli in the test trials, the practice trials also featured inharmonic stimuli).

**Figure 5.**
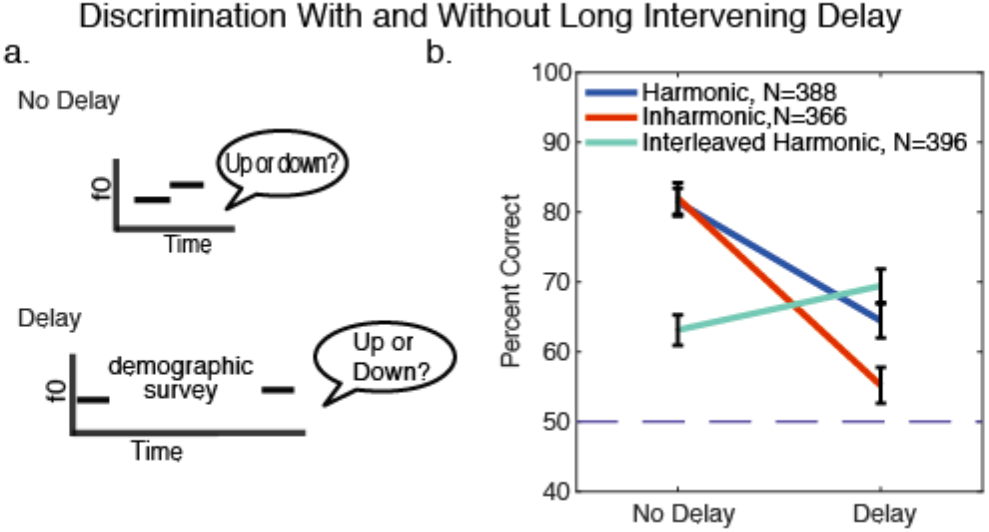
Experiment 5 – Harmonic advantage persists over longer delays with intervening task. a. Schematic of the two trial types in Experiment 5. Task was identical to that of Experiment 2. However, for the ‘delay’ condition participants were redirected to a short demographic survey that they could complete at their own pace. b. Results of Experiment 5 (which had the same stimulus conditions as Experiment 2). Error bars show standard error of the mean, calculated via bootstrap.

We ran a large number of participants to obtain sufficient power given the small number of trials per participant. Participants completed the survey at their own pace. We measured the time interval between the onset of the first note before the survey and the onset of the second note after the survey. Prior to analysis we removed participants who completed the survey in under 20 seconds (proceeding through the survey so rapidly as to suggest that the participant did not read the questions; 1 participant), or participants who took longer than 3 minutes (17 participants). Of the remaining 1150 participants, the mean time spent on the survey was 58.9 seconds (standard deviation of 25.1, median of 52.8 seconds and median absolute deviation of 18.4 seconds. There was no significant difference between the time taken for each of the conditions (Z<=0.94, p>=.345 for all pairwise comparisons).

As shown in Figure 5b, Experiment 5 qualitatively replicated the results of Experiments 1 and 2 even with the longer delay period and concurrent demographic survey. Without a delay, there was no difference between performance with harmonic and inharmonic conditions (p=.61, via bootstrap). With a delay, performance in both the Harmonic and Inharmonic conditions was worse than without a delay (p<.0001 for both). However, this impairment was larger for the Inharmonic condition; performance with inharmonic tones across a delay was significantly worse than that with harmonic tones (p<.001), producing an interaction between the type of tone and delay (p=.02). By contrast, performance on the Interleaved Harmonic condition did not deteriorate over the delay and in fact slightly improved (p=.004). This latter result could reflect the decay of the representation of the spectrum of the first tone across the delay, which in this condition might otherwise impair f0 discrimination by providing a competing cue (because the spectra of the two tones are different (28)).

### Experiment 6: Individual differences in tone discrimination

The similarity in performance between harmonic and inharmonic tones without a delay provides circumstantial evidence that listeners are computing changes in the same way for both types of stimuli, presumably using a representation of the spectrum in both cases. However, the results leave open the alternative possibility that listeners use a representation of the f0 for the harmonic tones despite having access to a representation of the spectrum (which they must use with the inharmonic tones), with the two strategies happening to support similar accuracy.

To address these possibilities, and to explore whether listeners use different encoding strategies depending on memory constraints, we employed an individual differences approach (45, 46). The underlying logic is that performance on tasks that rely on the same perceptual representations should be correlated across participants. For example, if two discrimination tasks rely on similar representations, a participant with a low threshold on one task should tend to have a low threshold on the other task.

In Experiment 6, we estimated participants’ discrimination thresholds either with or without a 3-second delay between stimuli, using a 3-down-1-up adaptive procedure (Figure 6a). Stimuli included harmonic, inharmonic, and interleaved-harmonic complex tones, generated as in Experiments 2-5. We used inharmonic stimuli for which a different pattern of jitter was chosen for each trial, because Experiment 3 showed similar results whether the inharmonic pattern was consistent or not, and because it seemed better to avoid a consistent jitter. Specifically, it seemed possible that fixing the jitter pattern across trials for each participant might create artifactual individual differences given that some jitter patterns are by chance closer to harmonic than others. Having the jitter pattern change from trial to trial produces a similar distribution of stimuli across participants and should average out the effects of idiosyncratic jitter patterns. We also included a condition with pure tones (sinusoidal tones, containing a single frequency) at the frequency of the fourth harmonic of the tones in the Harmonic condition, and a fifth condition where each note contained two randomly chosen harmonics from each successive set of four (1 to 4, 5 to 8, etc.). By chance, some harmonics could be found in both notes. This condition (Random Harmonic) was intended as an internal replication of the anticipated results with the Interleaved Harmonic condition.

**Figure 6.**
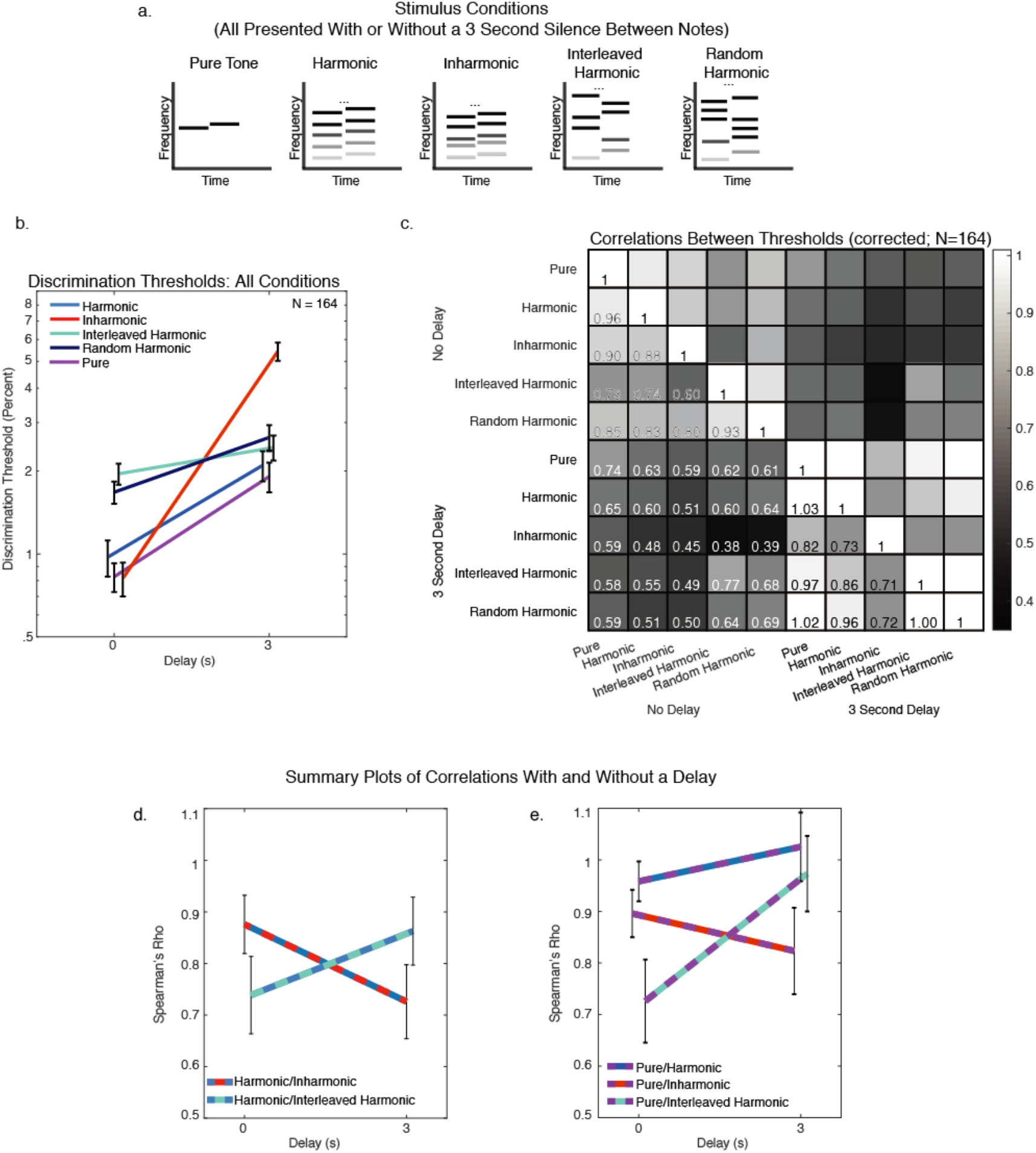
Experiment 6 – Individual differences suggest different representations depending on memory demands. a. Schematic of the five types of stimuli used in Experiment 6. Task and procedure (adaptive threshold measurements) were identical to those of Experiment 4, but with no inter-trial delays. b. Discrimination thresholds for all stimulus conditions, with and without a 3-second delay between tones. Here and in panels d-e, error bars show standard error of the mean, calculated via bootstrap. c. Matrix of the correlations between thresholds for all pairs of conditions. Correlations are Spearman’s rho, corrected for the reliability of the threshold measurements (i.e., corrected for attenuation). Corrected correlations can slightly exceed 1 given that they are corrected with imperfect estimates of the reliabilities. d. Comparison between Harmonic/Inharmonic and Harmonic/Interleaved Harmonic threshold correlations, with and without a delay. e. Comparison between Pure Tone condition correlation with and without a delay.

The main hypothesis we sought to test was that listeners use one of two different representations depending on the duration of the inter-stimulus delay and the nature of the stimulus. Specifically, without a delay, listeners use a detailed spectral representation for both harmonic and inharmonic sounds, relying on an f0-based representation only when a detailed spectral pattern is not informative (as in the Interleaved Harmonic condition). In the presence of a delay, they switch to relying on an f0-based representation for all harmonic sounds.

The key predictions of this hypothesis in terms of correlations between thresholds are 1) high correlations between Harmonic and Inharmonic discrimination without a delay, 2) lower correlations between Harmonic and Inharmonic discrimination with a delay, 3) low correlations between Interleaved Harmonic discrimination and both the Harmonic and Inharmonic conditions with no delay (because the former requires a representation of the f0), and 4) higher correlations between Interleaved Harmonic and Harmonic discrimination with a delay.

We ran this study online to recruit sufficient numbers to measure the correlations of interest. This experiment was relatively arduous (it took approximately 2 hours to complete), and based on pilot experiments we anticipated that many online participants would perform poorly relative to in-lab participants, perhaps because the chance of distraction occurring at some point over the 2 hours is high. To obtain a criterion level of performance with which to determine inclusion for online participants, we ran a group of participants in the lab to establish acceptable performance levels. We calculated the overall mean threshold for all conditions without a delay (five conditions) for the best two-thirds of in-lab participants, and excluded online participants whose mean threshold on the first run of those same 5 conditions (without a delay) was above this mean in-lab threshold. To avoid double dipping, we analyzed only the last three threshold measurements for each condition from online participants (10 total conditions – 3 runs for each condition). This inclusion procedure selected 164 of 450 participants for the final analyses shown in Figure 6b-e. See Supplementary Figure 5 for results with a less stringent inclusion criterion (the critical effects remained present with this less stringent criterion).

#### Mean Thresholds

The pattern of mean threshold measurements obtained online qualitatively replicated the results of Experiments 1-5 (Figure 6b). Inharmonic thresholds were indistinguishable from Harmonic thresholds without a delay (Z=1.74, p=.08), but were higher when there was a delay between sounds (Z=-9.07, p<.001). This produced a significant interaction between effects of tone type and delay (F(1,163)=88.45, p<.001, η_p_^2^=.35). And as in the previous experiments, there was no significant effect of delay for the Interleaved Harmonic condition (Interleaved Harmonic condition with vs. without delay; Z=0.35, p=.72).

#### Individual Differences – Harmonic, Inharmonic and Interleaved Harmonic conditions

Figure 6c shows the correlations across participants between different pairs of thresholds. Thresholds were correlated to some extent for all pairs of conditions, presumably reflecting general factors such as attention or motivation that produce variation in performance across participants. However, some correlations were higher than others. Figure 6d&e plot the correlations (extracted from the matrix in Figure 6c) for which our hypothesis makes critical predictions, to facilitate their inspection. Correlations here and elsewhere were corrected for the reliability of the underlying threshold measurements (47); the correlation between two thresholds was divided by the square-root of the product of their reliabilities (Cronbach’s alpha calculated from Spearman’s correlations between pairs of the last 3 runs of each condition). This denominator provides a ceiling for each correlation, as the correlation between two variables is limited by the accuracy with which each variable is measured. Thus, correlations could in principle be as high as 1 in the limit of large data, but because the threshold reliabilities were calculated from modest sample sizes, the corrected correlations could in practice slightly exceed 1.

Performance on Harmonic and Inharmonic conditions without a delay was highly correlated across participants (rho=.88, p<.001). However, the correlation between Harmonic and Inharmonic conditions with a delay was substantially lower (rho=.73, p<.001, significant difference between Harmonic-Inharmonic correlations with and without a delay, p=.007, calculated via bootstrap). We observed the opposite pattern for the Harmonic and the Interleaved Harmonic conditions: the correlation between Harmonic and Interleaved Harmonic thresholds was lower without a delay (rho=.74, p<.001) than with a delay (rho=.86, p<.001, significant difference between correlations, p=.03). This pattern of results yielded a significant interaction between the conditions being compared and the effect of delay (difference of differences between correlations with and without a delay = 0.27, p=.019. We replicated this interaction in a pilot version of the experiment that featured slightly different stimulus conditions and intervening notes in the delay period (Supplementary Figure 6; p=.006). Overall, the results of the individual differences analysis indicate that when listening to normal harmonic tones, participants switch between two different pitch mechanisms depending on the time delay between tones.

#### Results with Pure Tones and Random Harmonic Condition

The results for Pure Tones were similar to those for the Harmonic condition (Figure 6e), with quantitatively indistinguishable mean thresholds (without delay: Z=0.16, p=.87, with delay: Z=1.18, p=.24). Moreover, the correlations between Harmonic and Pure Tone thresholds were high both with and without a delay, and not significantly different (p=.13). These results suggest that similar representations are used to discriminate pure and harmonic complex tones. In addition, the correlations between the Pure Tone condition and the Inharmonic and Interleaved Harmonic conditions showed a similar pattern to that for the Harmonic condition (compare Figure 6e to Figure 6d), producing a significant interaction between the conditions being compared and the effect of delay (difference of differences between correlations with and without a delay = 0.32, p=.006). This interaction again replicated in the pilot version of the experiment (Supplementary Figure 6; p=.008).

The Random Harmonic results largely replicated the findings from the Interleaved Harmonic condition (Figure 6c). Evidently the changes that we introduced in the harmonic composition of the tones being compared in this condition were sufficient to preclude the use of the spectrum, and, as in the Interleaved Harmonic condition, performance was determined by f0-based pitch regardless of the delay between notes.

## Discussion

We examined the relationship between pitch perception and memory by measuring the discrimination of harmonic and inharmonic sounds with and without a time delay between stimuli. Across several experiments, we found that discrimination across a delay was better for harmonic sounds than for inharmonic sounds, despite comparable accuracy without a delay. This effect was observed over delays of a few seconds and persisted over longer delays with an intervening distractor task. We also analyzed individual differences in discrimination thresholds across a large number of participants. Harmonic and inharmonic discrimination thresholds were highly correlated without a delay between sounds, but were less correlated with a delay. By contrast, thresholds for harmonic tones and tones designed to isolate f0-based discrimination (interleaved-harmonic tones) showed the opposite pattern, becoming more correlated with a delay between sounds. Together, the results suggest that listeners use different representations depending on memory demands, comparing the spectra for sounds nearby in time, and the f0 for sounds separated in time. The results provide evidence for two distinct mechanisms for pitch discrimination, reveal the constraints that determine when they are used, and demonstrate a form of abstraction within the auditory system whereby the representations of memory differ in format from those used for rapid on-line judgments about sounds.

In hearing research, the word ‘pitch’ has traditionally referred to the perceptual correlate of the f0 (26). In some circumstances listeners must base behavior on the absolute f0 of a sound of interest, as when singing back a heard musical note. However, much of the time the information that matters to us is conveyed by how the f0 changes over time, and our results indicate that listeners often extract this information using a representation of the spectrum rather than the f0. One consequence of this is that note-to-note changes can be completely unambiguous even for inharmonic sounds that lack an unambiguous f0. Is this pitch perception? Under typical listening conditions (where sounds are harmonic) the changes in the spectrum convey changes in f0, and thus enable judgments about how the f0 changes from note to note. Consistent with this idea, listeners readily describe what they hear in the inharmonic conditions of our experiments as a pitch change, as though the consistent shift in the spectrum is interpreted as a change in the f0 even though neither note has a clear f0. We propose that these spectral judgments should be considered part of pitch perception, which we construe to be the set of computations that enable judgments about a sound’s f0. The perception of inharmonic “pitch changes” might thus be considered an illusion, exploiting the spectral pitch mechanism in conditions in which it does not normally operate.

Why don’t listeners base pitch judgments of harmonic sounds on their f0s when sounds are back-to-back? One possibility is that representations of f0 are in some cases less accurate and produce poorer discrimination than those of the spectrum. The results with interleaved harmonics (stimuli designed to isolate f0-based pitch) are consistent with this idea, as discrimination without a delay was worse for interleaved harmonics than for either harmonic or inharmonic tones that had similar spectral composition across notes. However, we note that this deficit could also reflect other stimulus differences, such as the potentially interfering effects of the changes in the spectrum from note to note (36). Regardless of the root cause for the reliance on spectral representations, the fact that performance with interleaved harmonics was similar with and without modest delays suggests that representations of the f0 are initially available in parallel with representations of the spectrum, with task demands determining which is used in the service of behavior. As time passes the high-fidelity representation of the spectrum appears to degrade, and listeners switch to more exclusively using a representation of the f0.

### Relation to previous studies of pitch and pitch memory

The use of time delays to study memory for frequency and/or pitch has a long tradition (48, 49). Our results here are broadly consistent with this previous work, but to our knowledge provide the first evidence for differences in how representations of the f0 and the spectrum are retained over time. A number of studies have examined memory for tones separated by various types of interfering stimuli, and collectively provide evidence that the f0 of a sound is retained in memory. For example, intervening sounds interfere most with discrimination if their f0s are similar to the tones being discriminated, irrespective of whether the intervening sounds are speech or synthetic tones (22), and irrespective of the spectral content of the intervening notes or the two comparison tones (21). Our results complement these findings by showing that memory for f0 has different characteristics from that for frequency content (spectra), by showing how these differences impact pitch perception, and by suggesting that memory for f0 should be viewed as a form of compression. We also show that introducing a delay between notes forces listeners to use the f0 rather than the spectrum, which may be useful in experimentally isolating f0-based pitch in future studies.

Our results are consistent with the idea that memory capacity limitations for complex spectra in some cases limit judgments about sound. Previous studies of memory for complex tones failed to find clear evidence for such capacity limitations, in that there was no interaction between the effects of inter-stimulus-interval and of the number of constituent frequencies on the accuracy of judgments of a remembered tone (18). However, there were many differences between these prior experiments and those described here that might explain the apparent discrepancy, including that the time intervals tested were short (at most 2 seconds) compared to those in our experiments, and the participants highly practiced. It is possible that under such conditions listeners are less dependent on the memory representations that were apparently tapped in our experiments. The tasks used in those prior experiments were also different (involving judgments of a single frequency component within an inharmonic complex tone), as were the stimuli (frequencies equidistant on a logarithmic scale). Quantitative models of memory representations and their use in behavioral tasks seem likely to be an important next step in evaluating whether the available results can be explained by a single type of memory store.

### Relation to visual memory

Our results could have interesting analogues in vision. Visual short-term memory has been argued to store relatively abstract representations (1-4, 6, 7), and the grouping of features into object-like representations is believed to increase its effective capacity (50-52). Our results raise the question of whether such benefits are specific to memory. It is plausible that for stimuli presented back-to-back, discrimination of visual element arrays would be similar irrespective of the element arrangement, with advantages for elements that are grouped into a coherent pattern only appearing when short-term memory is taxed. To our knowledge this has not been explicitly tested.

One apparent difference between auditory and visual memory is that in some contexts visual memory for simple stimuli decays relatively slowly, with performance largely unimpaired for multisecond delays comparable to those used here (24). By contrast, auditory memory is more vulnerable, with performance decreases often evident over seconds even for pure tone discrimination (e.g. Fig. 5b). As a consequence, visual memory has often been studied via memory ‘masking’ effects in which stimuli presented in the delay period impair performance if they are sufficiently similar to the stimuli being remembered (53-55). Such effects also occur for auditory memory (20-22), but the performance impairments that occur with a silent delay were sufficient in our case to illuminate the underlying representation. Masking effects might nonetheless yield additional insights.

### Relevance of f0-based pitch to music

F0-based pitch seems to be particularly important in music perception, evident in prior results documenting the effect of inharmonicity on music-related tasks. Melody recognition, ‘sour’ note detection, and pitch interval discrimination are all worse for inharmonic than for harmonic tones, in contrast to other tasks such as up/down discrimination, which can be performed equally well for the two types of tones (shown again in the experiments here) (31). Our results here provide a potential explanation for these effects. Music often requires notes to be compared across delays or intervening sounds, as when assessing a note’s relation to a tonal center (56), and the present results suggest that this should necessitate f0-based pitch. Musical pitch perception may have come to rely on representations of f0 as a result, such that even in musical tasks that do not involve storage across a delay, such as musical interval discrimination, listeners use f0-based pitch rather than the spectrum (31). It is possible that similar memory advantages occur for other patterns that occur frequently in music, such as common chords.

Given its evident role in music perception, it is natural to wonder whether f0-based pitch is honed by musical training. Western-trained musicians are known to have better pitch discrimination than Western non-musicians (57-59). However, previous studies examining effects of musicianship on pitch discrimination used either pure tone or harmonic complex tone stimuli, and thus do not differentiate between representations of the f0 vs. the spectrum. We found consistent overall pitch discrimination advantages for musicians compared to non-musicians (Supplementary Figures 1-3), but found no evidence that this benefit was specific to f0 representations: musicianship did not interact with the effects of inharmonicity or inter-stimulus delay. It is possible that more extreme variation in musical experience might show f0-specific effects. For instance, indigenous cultures in the Amazon appear to differ from Westerners in basic aspects of pitch (60) and harmony perception (61, 62), raising the possibility that they might also differ in the extent of reliance on f0-based pitch. It is also possible that musicianship effects might be more evident if memory were additionally taxed with intervening distractor tones.

### Efficient coding in perception and memory

Perception is often posited to estimate the distal causes in the world that generated a stimulus (63). Parameters that capture how a stimulus was generated are useful for behavior – as when one requires knowledge of an object’s shape in order to grasp it – but can also provide compressed representations of a stimulus. Indeed, efficient coding has been proposed as a way to estimate generative parameters of sensory signals (64). A sound’s f0 is one such generative parameter, and our results suggest that its representation may be understood in terms of efficient coding. Prior work has explained aspects of auditory representations (65, 66) and discrimination (67) as consequences of efficient coding, but has not explored links to memory. Our results raise the possibility that efficient coding may be particularly evident in sensory memory representations. We provide an example of abstract and compressed auditory memory representations, and in doing so explain some otherwise puzzling results in pitch perception (chiefly, the fact that conventional pitch discrimination tasks are not impaired by inharmonicity).

This efficient coding perspective suggests that harmonic sounds may be more easily remembered because they are prevalent in the environment, such that humans have acquired representational transforms to efficiently represent them (e.g. by projection onto harmonic templates) (68). This interpretation also raises the possibility that the effects described here might generalize to or interact with other sound properties. There are many other regularities of natural sounds that influence perceptual grouping (69, 70). Each of these could in principle produce memory benefits when sounds must be stored across delays. In addition to regularities like harmonicity that are common to a wide range natural sounds, humans also use learned ‘schemas’ for particular sources when segregating streams of sound (38, 69) and these might also produce memory benefits. It is thus possible that recurring inharmonic spectral patterns, for instance in inharmonic musical instruments (32), could confer a memory advantage to an individual with sufficient exposure to them, despite lacking the mathematical regularity of the harmonic series.

Memory could be particularly important in audition given that sound unfolds over time, with the structures that matter in speech, music, and other domains often extending over many seconds. Other examples of auditory representations that discard details in favor of more abstract structures include the ‘contour’ of melodies, that listeners retain in some conditions in favor of the exact f0 intervals between notes (71), or summary statistics of sound textures that average across temporal details (72, 73). These representations may reflect memory constraints involved in comparing two extended stimuli even without a pronounced inter-stimulus delay.

### Future directions

Our results leave open how the two representations implicated in pitch judgments are instantiated in the brain. Pitch-related brain responses measured in humans have generally not distinguished representations of the f0 from that of the spectrum (74-78), in part because of the coarse nature of human neuroscience methods. Moreover, we know little about how pitch representations are stored over time in order to mediate discrimination across a delay. In non-human primates there is evidence for representations of the spectrum of harmonic complex tones (79) as well as of their f0 (80), though there is increasing evidence for heterogeneity in pitch across species (81-85). Neurophysiological and behavioral experiments with delayed discrimination tasks in non-human animals could shed light on these issues.

Our results also indicate that we unconsciously switch between representations depending on the conditions in which we must make pitch judgments (i.e., whether there is a delay between sounds). One possibility is that sensory systems can assess the reliability of their representations and base decisions on the representation that is most reliable for a given context. Evidence weighting according to reliability is a common strategy in perceptual decisions (86), and our results raise the possibility that such frameworks could be applied to understand memory-driven perceptual decisions.

## Materials and Methods

### Participants

All experiments were approved by the Committee on the use of Humans as Experimental Subjects at the Massachusetts Institute of Technology, and were conducted with the informed consent of the participants. In all experiments except for Experiment 5 (which contained only two trials per participant), we excluded poorly performing participants using hypothesis-neutral performance criteria. In Experiments 1-4, we selected exclusion criteria a priori based on our expectations of what would constitute good performance. For Experiment 6, we conducted a pilot experiment in the lab, and selected online participants who performed comparably to good in-lab participants.

#### Musicianship

Across all experiments, musicians were defined as individuals with five or more years of self-reported musical training and/or active practice/performance. Non-musicians had four or fewer years of self-reported musical training and/or active practice/performance.

#### Experiment 1

16 participants were recruited for Experiment 1. 4 of these participants had an average performance across all conditions of less than 60% correct and their data were removed. The remaining participants (N=12, 5 female, mean age = 32.5 years, S.D. = 19.0 years) included 6 non-musicians and 6 musicians (24.5 years of training and active performance, S.D. = 16.0).

#### Experiment 2

34 participants were recruited for Experiment 2. 2 of these did not finish the experiment, and their data were not analyzed. Of the remaining 32 participants, 7 scored less than 60% correct across all conditions and were removed. The remaining participants (N=25, 12 female, mean age= 31.4 years, S.D.=14.7 years) included 10 non-musicians and 15 musicians (mean=16 years of musical training and active performance, S.D.=12.2 years).

#### Experiment 3

35 participants were recruited for Experiment 3. 3 of these did not complete the experiment or were removed for non-compliance. An additional 9 participants were removed from analysis because their average threshold across conditions (using the first run of each condition, allowing unbiased threshold estimates from the last three runs of each condition for the remaining participants) was greater than 3% (approximately half a semitone). The remaining participants (N=23, 9 female, mean age=36.0 years, S.D.=16.6 years) included 11 non-musicians and 12 musicians (mean=17.1 years of musical training and active, S.D.=13.5)

#### Experiment 4

77 participants were recruited online for Experiment 4. 39 participants were removed from analysis because their average threshold across the first run of all conditions was greater than 3% (approximately half a semitone). The high number of excluded participants is most likely due to the tedious nature of this experiment (because of the forced inter-trial interval in some of the conditions, which made it easy to lose focus). The remaining participants (N=38, 16 female, mean age=37.6 years, S.D.=11.4 years) included 27 non-musicians and 11 musicians (mean=8.5 years of musical training and active, S.D.=2.7).

#### Experiment 5

Before analysis, we removed the participants who completed the demographic survey in either less than 20 seconds, which we believed made it unlikely that those participants read and paid attention to all the questions, or greater than 3 minutes. 1 and 17 participants were excluded via these timing criteria, respectively. After these exclusions, 1150 people completed Experiment 5 (680 female, mean age = 35.0 years, S.D. = 11.6 years). 371 reported some form of musical training, and 290 of those reported greater than four years of training (mean=14.4 years, S.D.=9.2 years).

#### Experiment 6

450 participants were recruited for the online component of Experiment 6. We sought to obtain mean performance levels comparable with those of compliant and attentive participants run in the lab. To this end, we excluded participants whose average performance across conditions fell below a cutoff. We ran 10 participants in the lab to establish this cutoff. We used the average threshold from the best two-thirds of these in-lab participants (7 of the 10) across all 5 of the no-delay conditions as the cutoff for inclusion in the online experiment. This yielded a cutoff value of 2.18%, and 286 of the online participants were excluded from analysis because their average threshold (across all 5 of the no-delay conditions, using the first run for each of these conditions) was greater than this cutoff. We used only the first run to determine inclusion, and subsequently analyzed only the remaining three runs, to avoid selection bias in the threshold estimates. The remaining set of 164 participants (73 female, mean age=35.3 years, S.D.=9.1 years) included 73 who reported greater than four years of musical training (mean=11.0 years, S.D.=6.9 years).

### Audio Presentation: In-Lab

In all experiments, a MacMini computer running Psychtoolbox for MATLAB (87) was used to play sound waveforms. Sounds were presented to participants at 70 dB SPL over Sennheiser HD280 headphones (circumaural) in a soundproof booth (Industrial Acoustics). Sound levels were calibrated with headphones coupled to an artificial ear, with a microphone at the position of the eardrum. Participants logged their responses via keyboard press.

### Audio Presentation: Online

We used the crowdsourcing platform provided by Amazon Mechanical Turk to run experiments that necessitated large numbers of participants (Experiments 5 and 6), or when in-person data collection was not possible due to the COVID-19 virus (Experiment 4). Each participant in these studies used a calibration sound to set a comfortable level, and then had to pass a ‘headphone check’ experiment that helped ensure they were wearing headphones or earphones as instructed (44) before they could complete the full experiment. The experimental stimuli were set to 15.5 dB below the level of the calibration sound, to ensure that stimuli were never uncomfortably loud. Participants logged their responses by clicking buttons on their computer monitors using their mouse.

### Feedback

Feedback (correct/incorrect) was given after each trial for all tasks except for the two test trials for about half of the participants of Experiment 5 (see below).

### Experiment 1: Discriminating instrument notes with intervening silence

#### Procedure

Participants heard two instrument notes per trial, separated by varying amounts of silence (0, 5, and 10 seconds) and judged whether the second note was higher or lower than the first note. Participants heard 30 trials per condition, and all conditions were intermixed. The first stimulus for a trial began one second after the response was entered for the previous trial, such that there was at least a 1-second gap between successive trials.

#### Stimuli

Instrument notes were derived from the RWC Instrument database, which contains recordings of chromatic scales played on different instruments (88). We used recordings of baritone saxophone, cello, ukulele, pipe organ and oboe, chosen to cover a wide range of timbres. Instrument tones were manipulated using the STRAIGHT analysis and synthesis method (89-91). STRAIGHT is normally used to decompose speech into excitation and vocal tract filtering, but can also decompose a recording of an instrument into an excitation signal and a spectrotemporal filter. If the voiced part of the excitation is modeled sinusoidally, one can alter the frequencies of individual harmonics, and then recombine them with the unaltered instrument body filtering to generate inharmonic notes. This manipulation (89) leaves the spectral shape of the instrument largely intact. Previous studies with speech suggest that the intelligibility of inharmonic speech is comparable to that of harmonic speech (92). The frequency jitters for inharmonic instruments were chosen in the same way as the jitters for the inharmonic synthetic tones used in Experiments 2-6 (described below). The same pattern of jitter was used for both notes in a trial. STRAIGHT was also used to frequency-shift the instrument notes to create pairs of notes that differed in f0 by a specific amount. Audio was sampled at 16,000 Hz. All notes were 400 ms in duration and were windowed by 20 ms half-Hanning windows.

Each trial consisted of two notes. The second note differed from the first by .25 or .5 semitone. To generate individual trials, the f0 of the first note of each trial was randomly selected from a uniform distribution over the notes in a Western classical chromatic scale between 196 and 392 Hz (G3 to G4). A recording of this note, from an instrument selected from the set of 5 that were used (baritone saxophone, cello, ukulele, pipe organ and oboe), was chosen as the source for the first note in the trial (instruments were counterbalanced across conditions). If the second note in the trial was higher, the note 1 semitone above was used to generate the second note (the note 1 semitone lower was used if the second note of the trial was lower). The two notes were analyzed and modified using the STRAIGHT analysis and synthesis method (89-91); the notes were f0-flattened to remove any vibrato, shifted to ensure that the f0 differences would be exactly the intended f0 difference apart, and resynthesized with harmonic or inharmonic excitation. Some instruments, such as the ukulele, have slightly inharmonic spectra. These slight inharmonicities were removed for the Harmonic conditions due to the resynthesis.

### Stimuli for Experiments 2-6

Experiments 2, 3, 4, 5.1, and 6 used the same types of tones. The stimuli for Experiment 5.2 are described below. Synthetic complex tones were generated with exponentially decaying temporal envelopes (decay constant of 4 s^-1^) to which onset and offset ramps were applied (20 ms half-Hanning window). The sampling rate was 16,000 Hz for Experiment 2, and 48,000 Hz for all others. Prior to bandpass filtering, tones included all harmonics up to the Nyquist limit, in sine phase, and were always 400 ms in duration.

In order to make notes inharmonic, the frequency of each harmonic, excluding the fundamental, was perturbed (jittered) by an amount chosen randomly from a uniform distribution, *U*(-.5, .5). This jitter value was chosen to maximally perturb f0 (lesser jitter values did not fully remove peaks in the autocorrelation at the period of the original f0 (31)). Jitter values were multiplied by the f0 of the tone, and added to the frequency of the respective harmonic. For example, if the f0 was 200 Hz and a jitter value of −0.39 was selected for the second harmonic; its frequency would be set to 322 Hz. To minimize salient differences in beating, jitter values were constrained (via rejection sampling) such that adjacent harmonics were always separated by at least 30 Hz. The same jitter pattern was applied to every note of the stimulus for a given trial, such that the spectral pattern shifted coherently up or down, even in the absence of an f0. Except for the Inharmonic-Fixed condition of Experiment 3, where one random jitter pattern was used for entire blocks of the experiment, a new jitter pattern was chosen for each trial.

Each complex tone was band-pass filtered in the frequency domain with a Gaussian transfer function (in log frequency) centered at 2,500 Hz with a standard deviation of half an octave. This filter served to ensure that participants could not perform the tasks using changes in the spectral envelope, and also to minimize timbral differences between notes in the Interleaved Harmonic condition. The filter parameters were chosen to ensure that the f0 was attenuated (to eliminate variation in a spectral edge at the f0) while preserving audibility of resolved harmonics (harmonics below the 10^th^, approximately). The combination of the filter and the masking noise (described below) rendered the frequency component at the f0 inaudible.

To ensure that differences in performance for Harmonic and Inharmonic conditions could not be mediated by distortion products, we added masking noise to these bandpass filtered notes. We low pass filtered pink noise using a sigmoidal (logistic) transfer function in the frequency domain. The sigmoid had an inflection point at the third harmonic of the highest of the two notes on a trial, and a maximum slope yielding 40 dB of gain or attenuation per octave. We scaled the noise so that the noise power in a gammatone filter (one ERB_N_ in bandwidth (93), implemented as in (94)) centered at the f0 was 10 dB lower than the mean power of the three harmonics of the highest note of the trial that were closest to the 2,500 Hz peak (and thus had greatest magnitude) of the Gaussian spectral envelope (31). This noise power is sufficient to mask distortion products at the f0 (95, 96). This filtered and scaled pink noise was added to each note, and did not continue through the silence in ‘delay’ conditions. Noise has been reported to facilitate the perception of the f0 of a set of harmonics (97, 98) in contexts where the harmonic frequencies are embedded in relatively high levels of noise. Because the noise in our stimuli was focused at the f0 rather than the higher harmonics that composed our tones, it seems less likely to have produced such a benefit, but we never specifically manipulated it to assess its effect.

### Experiment 2: Discriminating synthetic tones with intervening silence

#### Procedure

The procedure was identical to that for Experiment 1, except stimuli were synthetic tones.

#### Stimuli

Each trial consisted of two notes, described above in **Stimuli for Experiments 2-6**. The second tone differed from the first by .25 or .5 semitones. The first note of each trial was randomly selected from a uniform distribution on a logarithmic scale spanning 200 to 400 Hz. Tones were either Harmonic, Inharmonic, or Interleaved Harmonic. Interleaved Harmonic notes were synthesized by removing harmonics [1,4, 5, 8, 9, etc.] in one note, and harmonics [2, 3, 6, 7, etc.] in the other note. They were otherwise identical to the Harmonic tones (identical bandpass filter in the frequency domain, as well as noise to mask a distortion product at the fundamental). This manipulation was intended to isolate f0-based pitch, as it removes the note-to-note spectral correspondence between harmonics.

### Experiment 3: Discriminating synthetic tones with a consistent inharmonic spectrum

#### Procedure

Participants heard two notes per trial, separated by varying amounts of silence (0, 1 and 3 seconds) and were asked whether the second note was higher or lower than the first note. Unlike Experiments 1 and 2, which used the method of constant stimuli, participants completed 2-up-1-down adaptive threshold measurements for each condition. Each run ended after 10 reversals. For the first 4 reversals, the f0 changed by a factor of 2, and for subsequent reversals by a factor of √2. Each adaptive run was initialized at an f0 difference of 1 semitone (approximately 6%), and the maximum f0 difference was limited to 16 semitones. The adaptive procedure continued if participants reached this 16 semitone limit; if they continued to get trials incorrect the f0 difference remained at the 16 semitone limit, and if they got two in a row right, the f0 difference would decrease from 16 semitones by a factor of 2 or √2 depending on how many reversals had already occurred. In practice participants who hit this limit repeatedly were removed before analysis due to our exclusion criteria. Thresholds were estimated by taking the geometric mean of the final 6 reversals. The first stimulus for a trial began one second after the response was entered for the previous trial, such that there was at least a 1-second gap between successive trials.

#### Stimuli

Each trial consisted of two notes, described above in **Stimuli for Experiments 2-6**. The first note of each trial was randomly selected from a uniform distribution on a logarithmic scale spanning 200 to 400Hz. Tones were either Harmonic, Inharmonic, or Inharmonic-Fixed, separated into three blocks, the order of which was counterbalanced across participants. Participants completed 12 adaptive thresholds within each block (3 delay conditions x 4 runs per condition). For the Inharmonic-Fixed block, a random jitter pattern was chosen at the beginning of the block and used for every trial within the entire block.

### Experiment 4: Discriminating synthetic tones with a longer inter-trial interval

#### Procedure

The procedure for Experiment 4 was identical to that for Experiment 3, except that the experiment was run on Amazon Mechanical Turk, with different time intervals between trials. Participants completed two sets of adaptive threshold measurements. In the first, trials were initiated by the participant, and could begin as soon as they entered the response for the previous trial and clicked a button to start the next trial. Four adaptive threshold measurements per condition were taken in this way (Harmonic and Inharmonic stimuli both with and without a 3-second delay). In the second set, a mandatory 4-second pause was inserted between each trial, which could be initiated by the participant once the pause had elapsed. Four threshold measurements for the same conditions were taken in this way, and thresholds were estimated by taking the geometric mean of the final 6 reversals. The two sets of measurements were randomly intermixed.

To perform adaptive procedures online, stimuli were pre-generated (instead of being generated in real-time as was done for the in-lab studies). For each condition and possible f0 difference, the stimuli were drawn randomly from a set of 20 pre-generated trials (varying in the f0 of the first note, and in the jitter pattern for the Inharmonic trials).

#### Stimuli

Stimuli were identical to those for the Harmonic and Inharmonic conditions in Experiment 3.

### Experiment 5: One-shot discrimination with longer intervening delay

#### Procedure

Participants were recruited using the Amazon Mechanical Turk crowdsourcing platform. In the main experiment, each participant completed two trials – one each of two trial types. In the first type of trial, they heard two consecutive notes and were asked whether the second note was higher or lower than the first. Notes always differed by a semitone. For the second type of trial, participants heard the first note, then were directed to a demographic survey, then were presented with the second note, and then were asked to respond. Before the presentation of the first note participants were told that they would be subsequently asked whether a second note heard after the survey was higher or lower in pitch than the note heard before the survey. Each participant heard the same stimulus type (Harmonic, Inharmonic, or Interleaved Harmonic) for each of the two trials. The order of the trials (with and without the intervening survey) was randomized across participants. For the trials with the demographic survey, we collected time stamps of when participants heard the first and second note in order to calculate the delay between notes. Because we thought feedback might affect the results, we ran two versions of the experiment. In the first version, participants received no feedback but were allowed to listen to the stimuli twice if they wished. In the second version there was feedback and participants heard each stimulus only once. 592 of the 1150 participants completed the first version of the experiment; the remaining 558 participants completed the second version. We found that there was no significant difference between the two versions of the experiment in any of the conditions. We thus combined data across the two versions.

Before completing the two main experiment trials and the survey, participants completed 10 practice trials without a delay, with the same type of tone that they would hear in the two main experiment trials (i.e. if a participant would hear inharmonic stimuli in the two experiment trials, their 10 practice trials would contain inharmonic stimuli). Participants received feedback on each of these practice trials. The stimulus difference in all trials was 1 semitone.

#### Stimuli

Stimuli for Experiment 5 were identical to those used in Experiments 2-4 (Harmonic, Inharmonic and Interleaved Harmonic conditions).

### Experiment 6: Individual differences in tone discrimination

#### Procedure

Both in lab and on Mechanical Turk, participants completed 2-up-1-down adaptive threshold measurements. The instructions were to judge whether the second note was higher or lower than the first note. The adaptive procedure was identical to that used in the first set of threshold measurements of Experiment 4 (each trial could be initiated by the participant as soon as they had entered their response for the previous trial). For the in-lab participants, we used the best three runs from each condition to set the inclusion criteria for online studies. For online participants, the first run of each condition was used to determine inclusion, and the final three runs of each condition were used for analysis. The order of the adaptive runs was randomized for each participant. There were 4 runs for each of the 10 conditions, for a total of 40 adaptive runs, randomized in order for each participant. Thresholds were estimated as the geometric mean of the final 6 reversals. Participants received feedback after each trial.

#### Stimuli

Participants in lab and online were tested on 5 different types of stimuli, presented either with no delay or a 3-second delay between tones (10 conditions). The five types of stimuli were as follows:

(1) Harmonic, (2) Inharmonic, and (3) Interleaved Harmonic, all identical to the same conditions in Experiment 2. (4) Pure Tones: We used the 4^th^ harmonic of the f0 (f0*4) so that the stimuli would overlap in frequency with the complex tones used in other conditions, which were filtered so that first audible harmonic was generally the 3^rd^ or 4^th^ harmonic. Low pass masking noise was omitted from the Pure Tone condition given that distortion products were not a concern. Given the similarity in mean thresholds between the Pure Tone and Harmonic conditions, and the high correlation between them across participants, the absence of noise in this condition does not appear to have influenced the results. (5) Random Harmonic: For each note, two harmonics were randomly chosen from harmonics 1 to 4, 5 to 8, etc. By chance, some harmonics could be present in both notes. This manipulation was intended to be intermediate between the Harmonic and Interleaved Harmonic conditions.

In all conditions, the f0 of the initial tone for each trial was chosen randomly from a uniform distribution on a logarithmic scale spanning 200-400 Hz.

As in Experiment 4, stimuli were pre-generated to enable online threshold measurements. For each condition and possible f0 difference, the stimuli were drawn randomly from a set of 20 pre-generated trials (varying in the f0 of the first note, and in the jitter pattern for the inharmonic conditions).

### Sample Sizes

#### Experiments 1 and 2

A power analysis of pilot data for Experiments 1 and 2 showed an effect size of d=1.25 for the difference between harmonic and inharmonic conditions at 5 seconds. We thus aimed to run at least 6 musicians and 6 non-musicians to be able to analyze the two groups separately and have an 80% chance of detecting the harmonic advantage at a p<.05 significance level (using paired t-tests). This number of participants also left us well-powered to observe an interaction between harmonicity and delay for both groups (8 participants were needed to have an 80% of detecting an interaction with an effect size of that seen in pilot data, η_p_^2^ = .2). Power analyses for Experiments 1-4 and Experiment 6 (in lab baseline) used G*Power (99). Experiment 2 was run in combination with other experiments (not described here) that were not as well powered and required more data, hence the additional participants.

#### Experiments 3 and 4

We performed power analyses for Experiments 3 and 4 using a pilot experiment with 17 participants where we measured thresholds either with or without a 3-second delay. The pilot experiment used the same method and analysis as Experiments 3 and 4, but without the 1-second-delay and Fixed-Jitter condition of Experiment 3 or the inter-trial delays of Experiment 4. The effect size from this pilot experiment for the Harmonic-Inharmonic difference at the 3-second delay was *d*=1.49. Based on the rough intuition that the effect of the Inharmonic-Fixed manipulation or the inter-trial delay might produce an effect approximately half this size, we sought to be 80% likely to detect an effect half as big as that observed in our pilot data, at a p<.05 significance (using a two-sided Wilcoxon signed-rank test). This yielded a target sample size of 17 participants. We did not plan to recruit equal numbers of musicians and non-musicians due to the similarity between groups in Experiments 1-2.

#### Experiment 5

For Experiment 5 we performed a power analysis by bootstrapping using a pilot version of the experiment (similar to the current version but without practice trials). For each possible sample size we computed the bootstrap distribution of the difference in performance between the Harmonic and Inharmonic conditions with a delay, as well as a null distribution obtained with conditions permuted across participants. We sought the sample size yielding an 80% chance of seeing a significant Harmonic-Inharmonic difference (i.e. where 95% of the bootstrap samples showed a difference exceeding the 97.5^th^ percentile of the null distribution), yielding a target sample size of 178 participants in each condition. We made no attempt to recruit equal numbers of musicians and non-musicians, as we did not plan to analyze those groups separately given the similar results across groups observed in the experiments we ran prior to this.

#### Experiment 6

For Experiment 6, we performed a power analysis by bootstrapping pilot data (an earlier version of Experiment 6 with slightly different stimuli). For each of a set of sample sizes we computed bootstrap distributions of the interaction term (difference of differences between the conditions being compared, Harmonic/Inharmonic and Interleaved Harmonic), as well as null distributions obtained by permuting conditions across participants. We found that a sample size of 154 yielded a 90% chance of seeing the interaction present in our pilot data at a p<.05 significance level. We ran more than this number of participants to allow performance-based exclusion. As with Experiments 3-5, we made no attempt to recruit equal numbers of musicians and non-musicians.

### Statistics

For Experiments 1, 2, and 5 we calculated percent correct for each condition. For Experiments 1 and 2, data were evaluated for normality with Lilliefors’ composite goodness-of-fit test. Data for Experiment 1 passed Lilliefors’ test, and so significance was evaluated using paired t-tests and repeated-measures ANOVAs. We used mixed-model ANOVAs to examine the effects of musicianship (to compare within- and between-group effects). Data for Experiment 2 were nonnormal due to ceiling effects in some conditions, and so significance was evaluated with the same non-parametric tests used for the threshold experiments (described below).

For Experiment 5, the significance of the differences between conditions and the significance of interactions were calculated via bootstrap (10,000 samples). To calculate the significance of the interaction between conditions, we first calculated the interaction (the difference of differences in means with and without a delay). For instance, for the Harmonic and Inharmonic conditions this term is as follows:

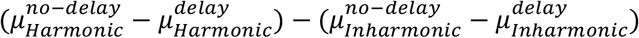

Then, we approximated a null distribution for this interaction, permuting conditions across participants and recalculating the difference of differences 10,000 times. To determine statistical significance we compared the actual value of the interaction to this null distribution.

Data distributions were non-normal (skewed) for threshold experiments (Experiments 3, 4 and 6) as well as for Experiment 2, so non-parametric tests were used for all comparisons. Wilcoxon signed-rank tests were used for pairwise comparisons between dependent samples (for example, two conditions for the same participant group). To compare performance across multiple conditions or across musicianship we used F statistics for repeated-measures ANOVAs (for within group effects) and mixed-model ANOVAs (to compare within and between group effects). However, because data were non-normal, we evaluated the significance of the F statistic with approximate permutation tests, randomizing the assignment of the data points across the conditions being tested 10,000 times, and comparing the F statistic to this distribution.

We used Spearman’s rank correlations to examine individual differences in Experiment 6. Correlations were corrected for the reliability of the threshold measurements using the Spearman correction for attenuation (47). We used standardized Cronbach’s alpha as a measure of reliability (100, 101). This entailed calculating the Spearman correlation between pairs of the 3 analyzed adaptive track thresholds for each condition, averaging these three correlations, and applying the Spearman-Brown correction to estimate the reliability of the mean of the three adaptive threshold measurements. Standard errors for correlations were estimated by bootstrapping the correlations 10,000 times. To calculate the significance of the interaction between conditions, we first calculated the interaction (the difference of differences). For instance, for the Harmonic and Inharmonic conditions, this term was:

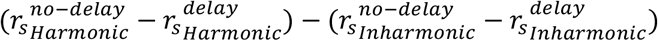

Then, we approximated a null distribution for this interaction, permuting conditions across participants and recalculating the difference of differences 10,000 times. To determine statistical significance we compared the actual value of the interaction to this null distribution.

## Acknowledgements

The authors thank R. Grace, S. Dolan, and C. Wang for assistance with data collection, T. Brady for helpful discussions, and L. Demany and the entire McDermott Lab for comments on the manuscript. The work was supported by the McDonnell Foundation Scholar Award and NIH grant No. R01DC014739 to J.H.M. and NIH grant F31DCO18433 and a NSF Graduate Research Fellowship to M.J.M.

## Author Contributions

M.J.M. designed the experiments, collected and analyzed the data, made the figures, and wrote the paper. J.H.M. designed the experiments and wrote the paper.

## Supplementary Figures

**Supplementary Figure 1:**
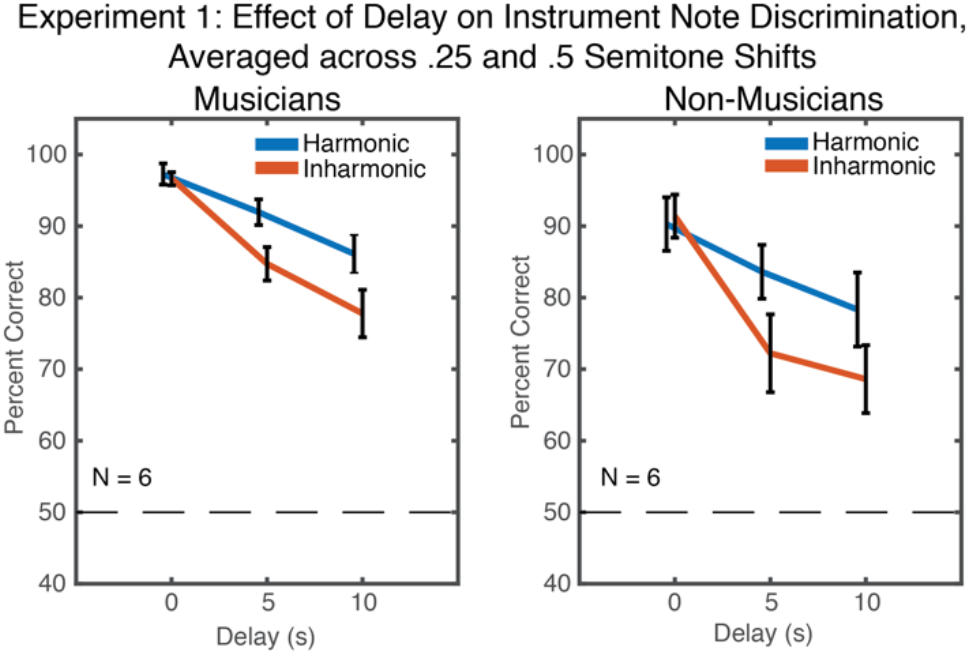
Results from Experiment 1, plotted separately for musicians and non-musicians. Results are averaged across the two difficulty levels (.25 and .5 semitones) to maximize power. Error bars show standard error of the mean. There was no interaction between musicianship, harmonicity and delay length (F(2,20)=0.58, p=.57, η_p_^2^=.06), and the interaction between delay and harmonicity was significant in non-musicians alone (F(2,10)=6.48, p=.02, η_p_^2^=.56)).

**Supplementary Figure 2.**
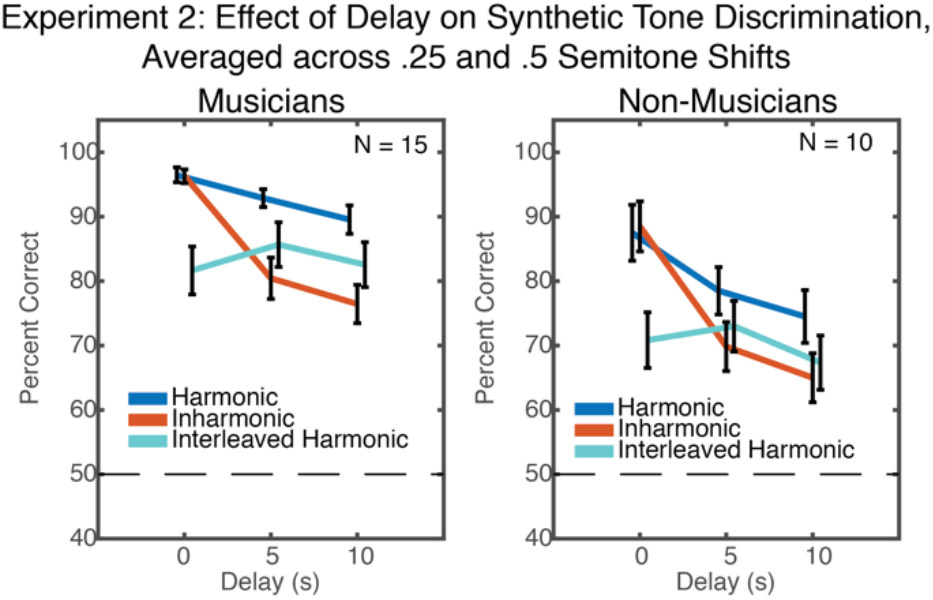
Results from Experiment 2, plotted separately for musicians and non-musicians. Results are averaged across the two difficulty levels (.25 and .5 semitones) to maximize power. Error bars show standard error of the mean. As in Experiment 1, the effects were qualitatively similar for musicians and non-musicians. Although there was a significant main effect of musicianship (F(1,23)=10.28, p<.001, η_p_^2^=.99), the interaction between the effects of delay and harmonicity was significant in both musicians (F(2,28)=20.44, p<.001, η_p_^2^=.59) and non-musicians (F(2,18)=11.99, p<.001, η_p_^2^=.57), and there was no interaction between musicianship, stimulus type (Harmonic, Inharmonic, Interleaved Harmonic), and delay length (F(4,92)=0.19, p=.98, η_p_^2^=.01).

**Supplementary Figure 3.**
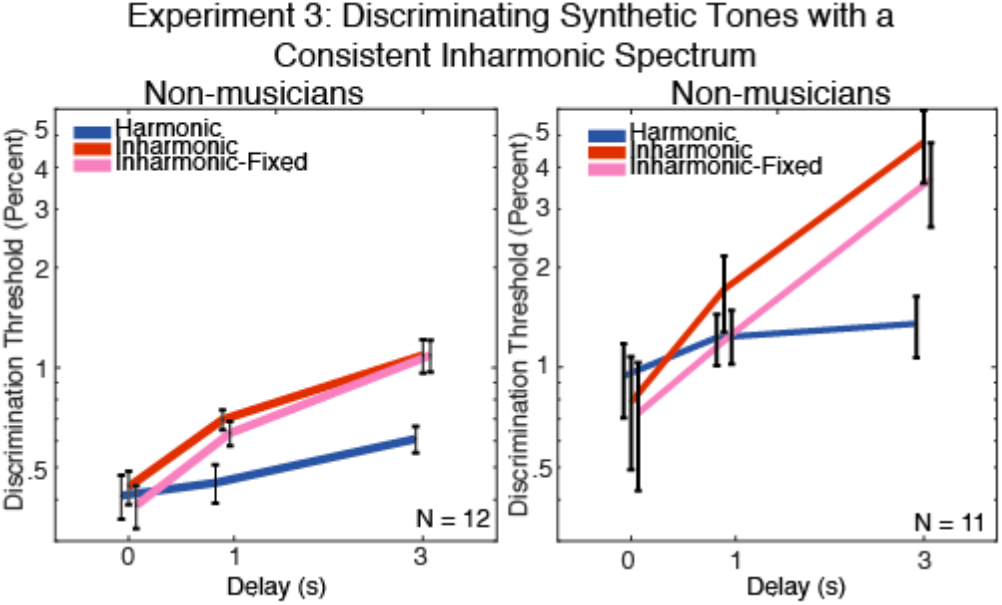
Results from Experiment 3, plotted separately for musicians and non-musicians. Error bars show within-subject standard error of the mean. We again observed significant interactions between the effects of delay and harmonicity in both musicians (F(4,44)=3.85, p=.009, η_p_^2^=.26) and non-musicians (F(4,40)=3.04, p=.028, η_p_^2^=.23), and no interaction between musicianship, stimulus type (Harmonic, Inharmonic, Inharmonic-Fixed), and delay length (F(4,84)=1.05, p=.07, η_p_^2^=.05).

**Supplementary Figure 4.**
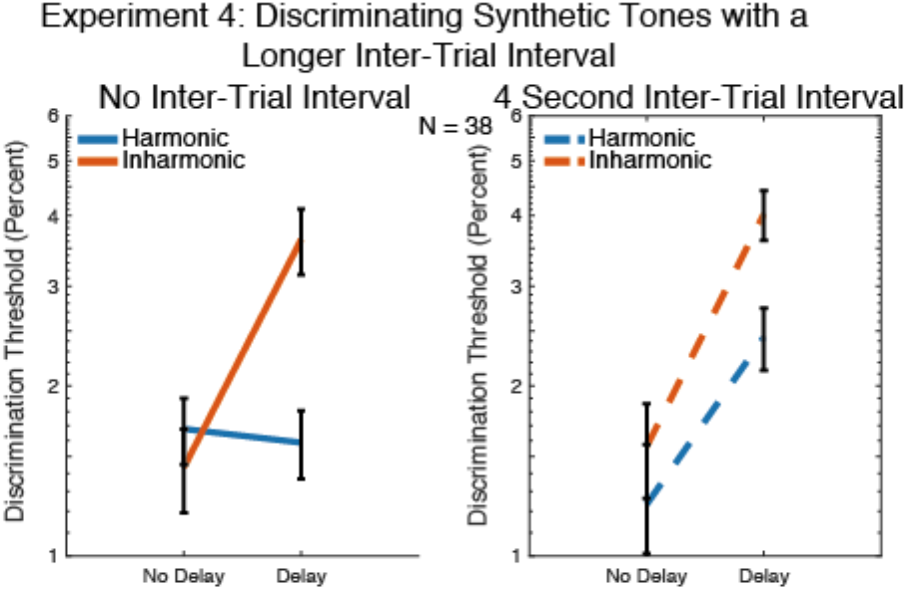
Results from Experiment 4, measuring discrimination of synthetic tones with and without a delay between notes, and with and without a longer inter-trial interval. Results from trials with (right) and without (left) an added 4 second delay between trials are plotted separately. Error bars show within-subject standard error of the mean. The interaction between within-trial delay (0 vs. 3 seconds) and stimulus type (Harmonic vs. Inharmonic) was present both with and without the longer inter-trial interval (with: F(1,37)=4.92, p=.03, η_p_^2^=.12; without: F(1,37)=12.34, p=.001, η_p_^2^=.25).

**Supplementary Figure 5.**
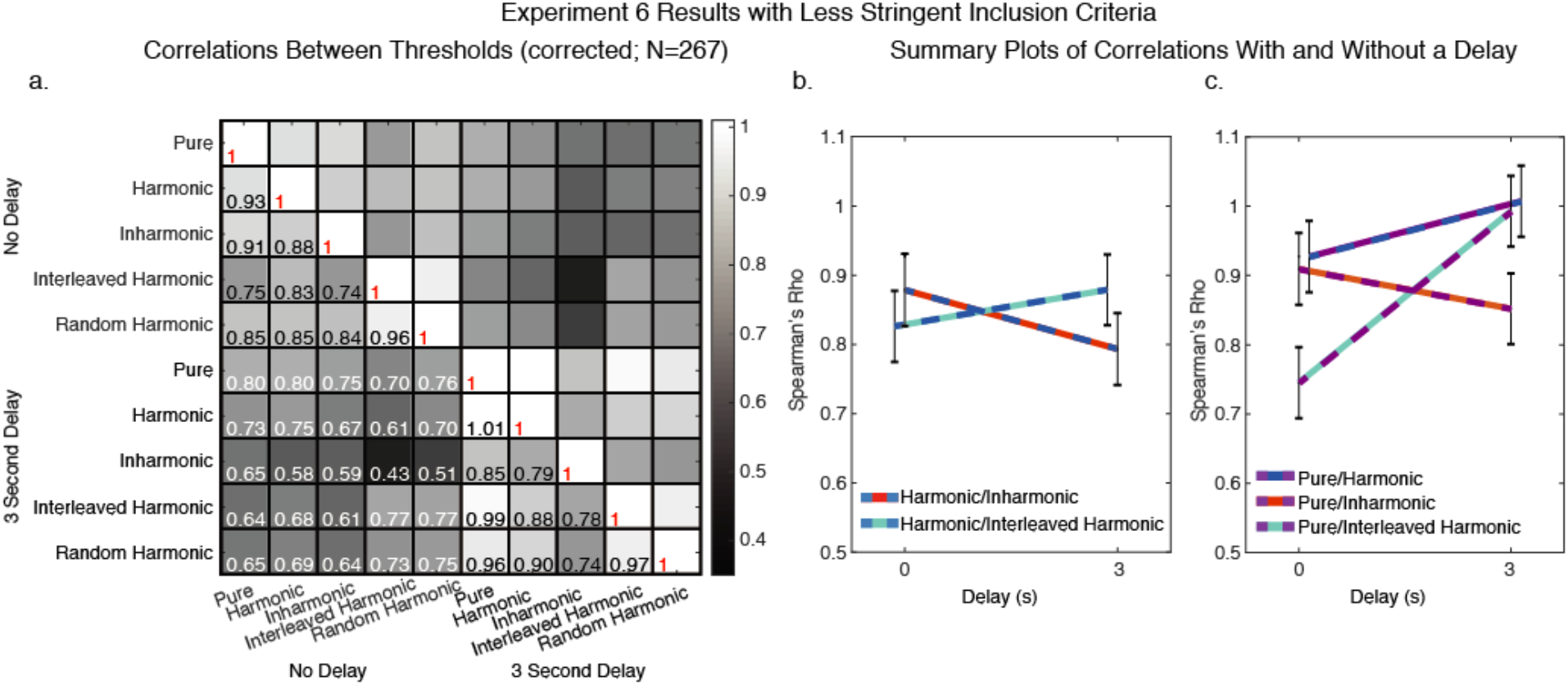
Individual Differences Results with Less Stringent Inclusion Criteria. Instead of including only those participants who performed as well as in-lab participants, we excluded participants if their average threshold across all 5 conditions on the first run of the no-delay trials was greater than 5% (just under 1 semitone). This excluded 183 of 450 participants, leaving 267 participants (136 female, mean age=35.8 years, S.D.=9.5 years). 94 of these participants reported greater than four years of musical training (mean=10.8, S.D.=6.6 years). (A) Matrix of the correlation between thresholds for all pairs of conditions. Correlations are Spearman’s rho, corrected for the reliability of the threshold measurements (i.e., corrected for attenuation). (B) Comparison between Harmonic/Inharmonic and Harmonic/Interleaved Harmonic correlations, with and without a delay. The interaction between Harmonic/Inharmonic and Harmonic/Interleaved Harmonic correlations remained significant even with this more lenient inclusion criteria (difference of differences between correlations with and without a delay = 0.14, p=.044). Error bars in b and c show standard error of the mean, calculated via bootstrap. (C) Comparison between Harmonic/Pure, Inharmonic/Pure, and Interleaved Harmonic/Pure correlations, with and without a delay. The interaction between the Inharmonic/Pure and Interleaved Harmonic/Pure correlations likewise remained significant with the less stringent inclusion criteria (difference of differences between correlations with and without a delay = 0.30, p<.001).

**Supplementary Figure 6.**
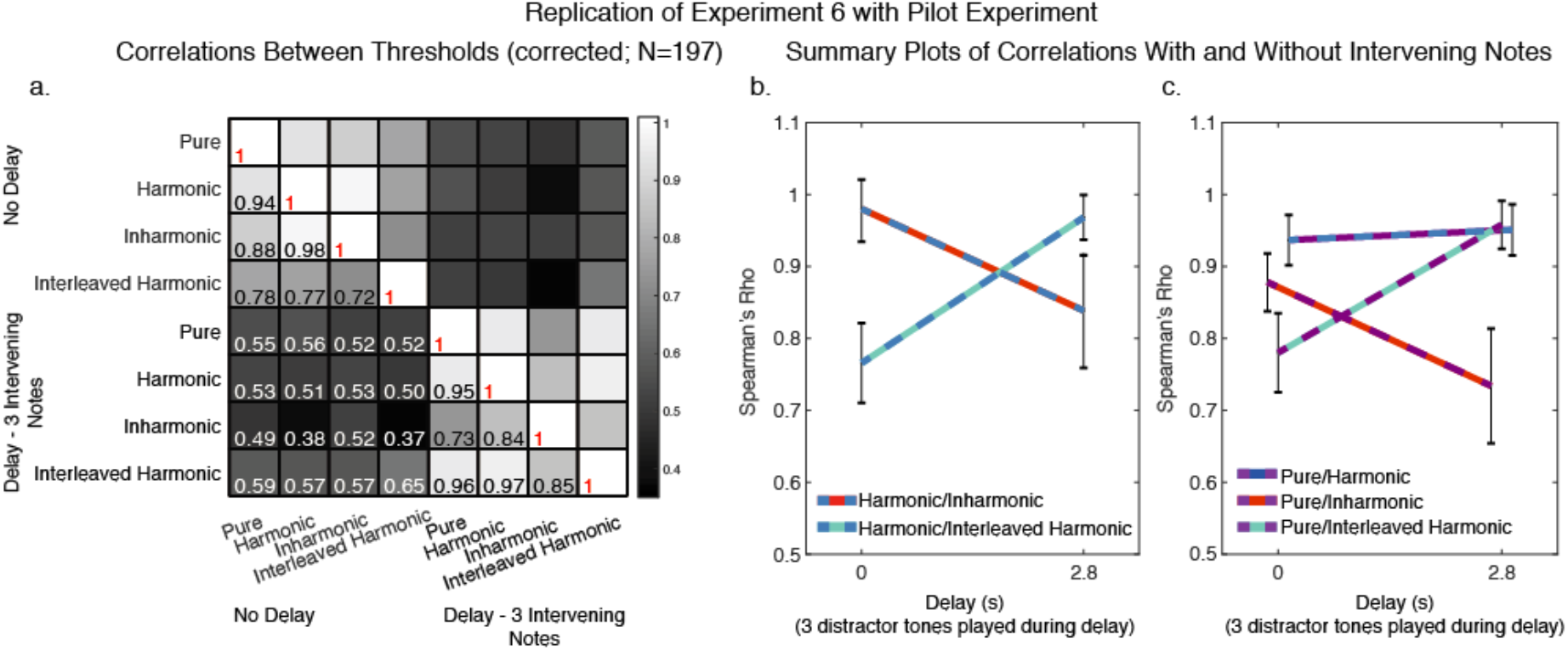
Replication of Individual Differences Results with Pilot Experiment. Results of Experiment 6 were replicated in two pilot experiments, the data from which are combined in this figure. Both pilot experiments were run online using 2-down-1-up adaptive procedures. Participants discriminated tones identical to those used in Experiment 6, except for the Random Harmonic conditions, which we thus omitted from this figure. Instead of having a 3-second silent delay between test tones, both pilot experiments presented three intervening distractor notes between the test tones. In these conditions, participants heard the first test tone, a 200ms pause, three back-to-back 800ms notes, a 200ms pause, and then the second test tone (yielding a total delay between the two test tones of 2.8 seconds). For two of the four adaptive runs, intervening notes were harmonic, and for the other two runs they were inharmonic (intervening tones were generated in the same way as the main test tones). The runs for all stimulus conditions were randomly ordered throughout the experiment. 310 participants completed the first pilot experiment, in which the intervening notes were chosen randomly from a 7-semitone distribution surrounding the first note (loosely modeled after the method used in Semal & Demany, 1990). 295 participants completed the second pilot experiment, in which the 3 intervening notes were chosen randomly from a uniform distribution spanning 178.2 Hz-449 Hz (200-400 Hz +/- 2 semitones). In the first pilot experiment, adaptive runs for tones without an inter-stimulus delay (and thus without intervening notes) were initialized at 1 semitone pitch difference, and adaptive tracks for intervening note conditions were initialized at a 2 semitone pitch difference. For the second pilot experiment, all adaptive tracks were initialized at a 2 semitone pitch difference. Because results from the two pilots were similar, we combined the data, and then used the same filtering procedure used in Experiment 6 - participants who performed worse than 2.18% across the first (of four) runs on conditions without intervening notes were removed from further analysis. This excluded 408 of the total 605 participants, leaving 197 participants (77 female, mean age=34.2 years, S.D.=9.8 years). 52 of these participants reported greater than four years of musical training (mean=10.9, S.D.=8.8 years). (A) Matrix of the correlation between thresholds for all pairs of conditions. Correlations are Spearman’s rho, corrected for the reliability of the threshold measurements (i.e., corrected for attenuation). (B) Comparison between Harmonic/Inharmonic and Harmonic/Interleaved Harmonic threshold correlations, with and without a delay. The interaction between Harmonic/Inharmonic and Harmonic/Interleaved Harmonic correlations was significant in this pilot study (difference of differences between correlations with and without a delay = 0.34, p=.006), replicating the effect from Experiment 6. Error bars show standard error of the mean, calculated via bootstrap. (C) Comparison between Harmonic/Pure, Inharmonic/Pure, and Interleaved Harmonic/Pure correlations, with and without a delay. The interaction between the Inharmonic/Pure and Interleaved Harmonic/Pure condition was also significant (difference of differences between correlations with and without a delay = 0.32, p=.008), again replicating the effect seen in Experiment 6.

